# Derivation of human trophoblast stem cells from placentas at birth

**DOI:** 10.1101/2024.05.01.592064

**Authors:** Victoria Karakis, John W. Britt, Mahe Jabeen, Adriana San Miguel, Balaji M Rao

**Author notes:** These authors contributed equally (Victoria Karakis, John W. Britt, Mahe Jabeen). **Corresponding author**: Dr. Balaji Rao.

## Abstract

Human trophoblast stem cells (hTSCs) have emerged as a powerful tool for modeling the placental cytotrophoblast (CTB) in vitro. hTSCs were originally derived from CTBs of the first trimester placenta or blastocyst-stage embryos in trophoblast stem cell medium (TSCM) that contains epidermal growth factor (EGF), the glycogen synthase kinase-beta (GSK3β) inhibitor CHIR99021, the transforming growth factor-beta (TGFβ) inhibitors A83-01 and SB431542, valproic acid (VPA), and the Rho-associated protein kinase (ROCK) inhibitor Y-27632. Here we show that hTSCs can be derived from CTBs isolated from the term placenta, using TSCM supplemented with a low concentration of mitochondrial pyruvate uptake inhibitor UK5099 and lipid-rich albumin (TUA medium). Notably, hTSCs could not be derived from term CTBs using TSCM alone, or in the absence of either UK5099 or lipid-rich albumin. Strikingly, hTSCs cultured in TUA medium for a few passages could be transitioned into TSCM and cultured thereafter in TSCM. hTSCs from term CTBs cultured in TUA medium as well as those transitioned into and cultured in TSCM thereafter could be differentiated to the extravillous trophoblast and syncytiotrophoblast lineages and exhibited high transcriptome similarity with hTSCs derived from first trimester CTBs. We anticipate that these results will enable facile derivation of hTSCs from normal and pathological placentas at birth with diverse genetic backgrounds and facilitate in vitro mechanistic studies in trophoblast biology.

**Significance statement:** Human trophoblast stem cells (hTSCs) derived from early pregnancy have emerged as a powerful tool for modeling early human placental development in vitro. However, restrictions on derivation of cell lines from fetal tissues limit the availability and genetic diversity of hTSC lines. Also, the associated pregnancy outcome of hTSCs derived from early gestation is unknown. Here we describe the derivation and characterization of hTSCs from placentas obtained at birth. Our results will enable derivation of genetically diverse hTSCs associated with normal and pathological pregnancies, enabling research in placental biology.

## Introduction

Upon embryo implantation, the outer trophectoderm layer of the blastocyst-stage embryo gives rise to the epithelial cytotrophoblast (CTB), which forms the two major differentiated cell lineages in the placenta – the multinucleate syncytiotrophoblast (STB) and the extravillous trophoblast (EVT) (1). Dysfunctional trophoblast development leads to defects in placentation and contributes to several pregnancy-related pathologies including pre-eclampsia, intrauterine growth restriction, and preterm birth (2, 3). However, mechanistic studies on early human trophoblast development are challenging due to restrictions on research with human embryos and fetal tissue, limited availability of placental samples from early gestation, and significant differences in placental development in humans and other commonly used animal models (4). Therefore, there is significant interest in the development of in vitro models that accurately mimic human trophoblast development in vivo.

In this context, human trophoblast stem cells (hTSCs) – first derived by Okae et al. (5) from first trimester CTBs and blastocyst-stage embryos – have emerged as a powerful in vitro model for studying human trophoblast development. Analogous to the placental CTB, hTSCs can be expanded in cell culture, and can be differentiated to the STB and EVT lineages. In other studies, Haider et al. (6) and Turco et al. (7) derived self-renewing trophoblast organoids (TO) from first trimester CTBs. Unlike hTSCs that are relatively homogenous in 2D culture, TOs are 3D structures that contain both CTB-like multipotent cells and STB.

Arguably, a limitation of hTSCs and TOs derived from first trimester placental tissues, obtained through elective termination of pregnancy, is that the associated pregnancy outcome is typically unknown. As an alternative, several studies have established hTSCs by differentiation of human embryonic stem cells (hESCs) derived from the inner cell mass of blastocyst-stage embryos or human induced pluripotent stem cells (hiPSCs) obtained by reprogramming of somatic cells by overexpression of pluripotency-associated genes (8–10). Additionally, somatic cells have been directly reprogrammed to form human induced trophoblast stem cells (hiTSCs) by overexpression of trophoblast-specific genes (11, 12). These approaches potentially enable the development of hTSC models using somatic cells associated with known normal or pathological outcomes. However, further work is needed to assess epigenetic differences between hTSC models obtained from reprogrammed somatic cells and primary placental tissue.

Placental growth continues throughout gestation, suggesting that trophoblast stem cells may be present even in placentas at term. Yet, hTSCs or TOs cannot be derived from term placentas using previously described protocols (5–7). Nevertheless, results from two recent studies suggest that hTSCs and TOs can indeed be derived from term placentas. Yang et al. modified previous protocols described by Turco et al. (7) to derive organoids from term placentas (13). Wang et al. have derived hTSCs from term placentas by culturing CTBs in trophoblast stem cell medium (TSCM) described by Okae et al., at 1% oxygen (14).

In this study, we report the derivation and characterization of hTSCs using CTBs isolated from term placentas, in TSCM supplemented with a low concentration (50 nM) of the mitochondrial pyruvate uptake inhibitor UK5099 and AlbuMAX II, i.e. lipid-rich albumin (TUA medium). Notably, our culture conditions do not require the use of hypoxic conditions. Furthermore, we show that cells cultured for a few passages in TUA medium can be transitioned to TSCM and cultured further, like hTSCs derived from first trimester placentas. hTSCs from term placentas derived and maintained in TUA medium, as well as those transitioned into TSCM, exhibit high transcriptome similarity with hTSCs from first trimester placentas and can be differentiated to EVT and STB in vitro. The ability to easily generate hTSC lines from normal and pathological placentas at birth will accelerate mechanistic studies in human placental biology.

## Results

### Derivation of hTSCs from term CTBs in TUA medium

Okae et al. derived hTSCs from first trimester placentas and blastocyst-stage embryos in TSCM that contains epidermal growth factor (EGF), the glycogen synthase kinase-beta (GSK3β) inhibitor CHIR99021, the transforming growth factor-beta (TGFβ) inhibitors A83-01 and SB431542, valproic acid (VPA), and the Rho-associated protein kinase (ROCK) inhibitor Y27632. We derived two hTSC lines (T1(female) and T2 (male)) using CTBs isolated from term placentas with no known pathology using TSCM supplemented with a low concentration (50 nM) of the mitochondrial pyruvate uptake inhibitor UK5099 and lipid-rich bovine serum albumin (AlbuMAX II, 0.2%); this medium is referred to as TUA medium. Briefly, CTBs from term placentas expressing CD49f (integrin α6) were isolated as described in Materials and Methods and cultured in TUA medium on tissue culture plates coated with vitronectin/laminin-521. Similar to observations by Okae et al.(5), we observed slow growth initially; however, highly proliferative cells with morphology similar to hTSCs derived from first trimester placentas were seen within a few passages (**Figs. 1A**). Under these conditions, cells express the pan-trophoblast marker KRT7, and markers associated with hTSCs including GATA3, TFAP2C, YAP, TEAD4, and p63 (**Figs. 1A, S1A**). Data from CT29 and CT30 hTSCs derived from first trimester placentas is shown for comparison in **Fig. S1B-D**. Interestingly, in contrast to CT29 and CT30 hTSCs, T1 and T2 hTSCs from term placentas showed expression of the early trophoblast marker CDX2 (**Figs. 1A, S1A, C-D**). Quantitative comparisons of CDX2 and p63 expression intensity in immunofluorescence images are shown in **Figs. 1B, S1B**. T1 and T2 hTSCs from term placentas have been maintained for 20+ passages in TUA medium, indicating robustness of these culture conditions.

**Figure 1:**
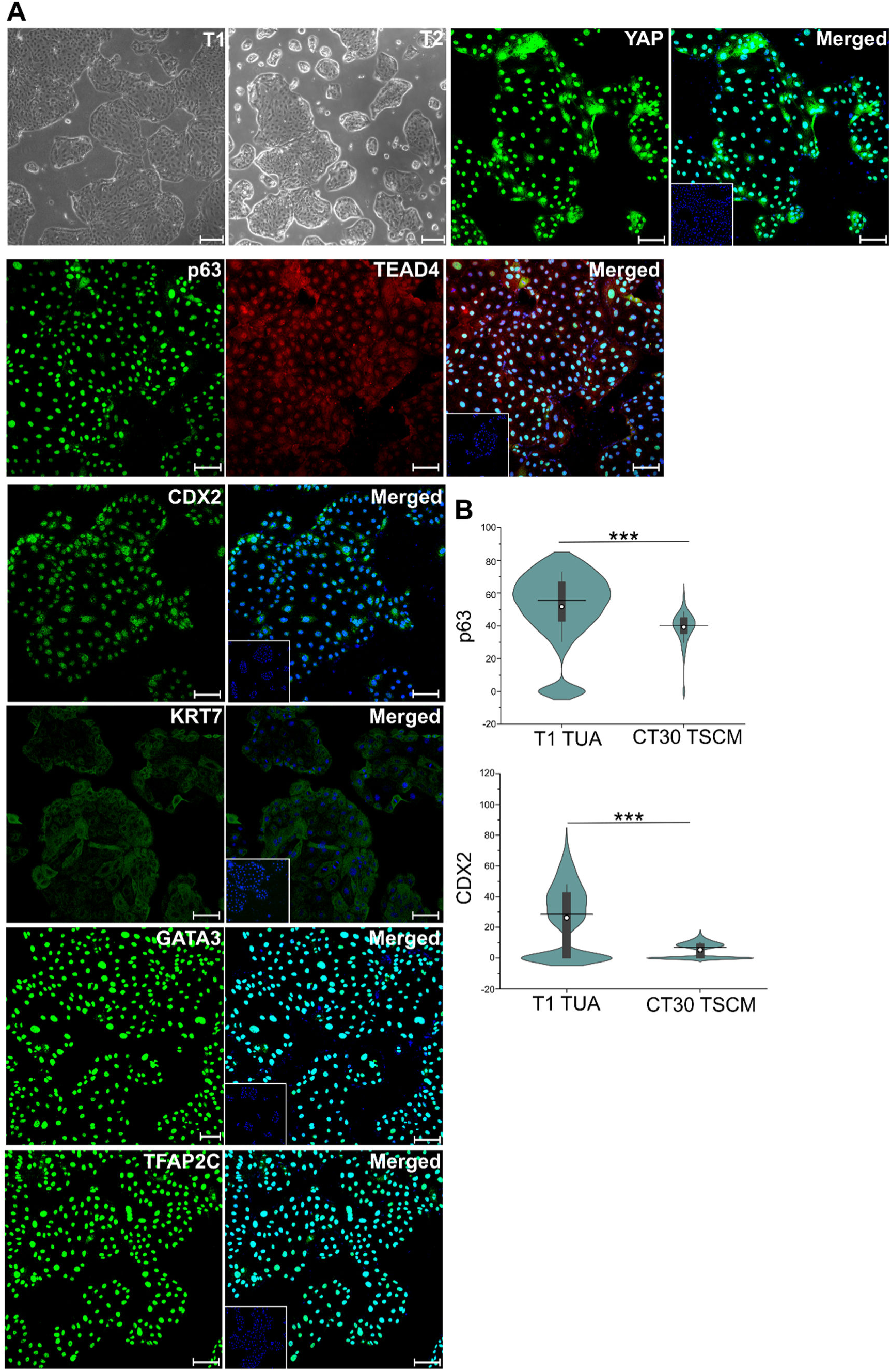
Expression of hTSC markers in TUA medium. (A) Bright field images for T1 and T2 hTSCs in TUA medium. Confocal microscopy imaging of T1 hTSCs cultured in TUA medium, staining for TFAP2C, YAP, GATA3, KRT7, TEAD4, p63, and CDX2; p63 was co-stained with TEAD4. Nuclei were stained with DAPI (blue). Isotype control is shown as an inset image. Scale bars are 100 µm for confocal images and 250 μm for bright field images. (B) Quantification of expression of p63 and CDX2 from T1 hTSCs (n= 5565 for p63, n= 6631 for CDX2) and CT30s hTSCs (n=1259 for p63, n=7813 for CDX2). Data from two biological replicates used. White circle represents the mean and the black line represents the median (***p-value < 0.001)

We further assessed the differentiation of T1 and T2 hTSCs derived from term CTBs to the STB and EVT lineages using chemically defined differentiation conditions previously described by Karakis et al. (15), or using protocols described by Okae et al (5) (**Figs. S2A, D**). Differentiation of CT29 and CT30 hTSCs derived from first trimester placentas was used as a comparison. (**Fig. S3**) With both protocols for EVT differentiation, T1 and T2 hTSCs gave rise to cells expressing the EVT markers NOTCH1 and HLA-G (**Figs. 2A**, **S2B**). Quantitative image analysis showed that protein expression levels of NOTCH1 and HLA-G were comparable between EVTs from T1 and T2 hTSCs from term placentas and CT29 and CT30 hTSCs previously derived by Okae et al. under both sets of differentiation conditions used (**Figs. 2B, S2C**).

**Figure 2:**
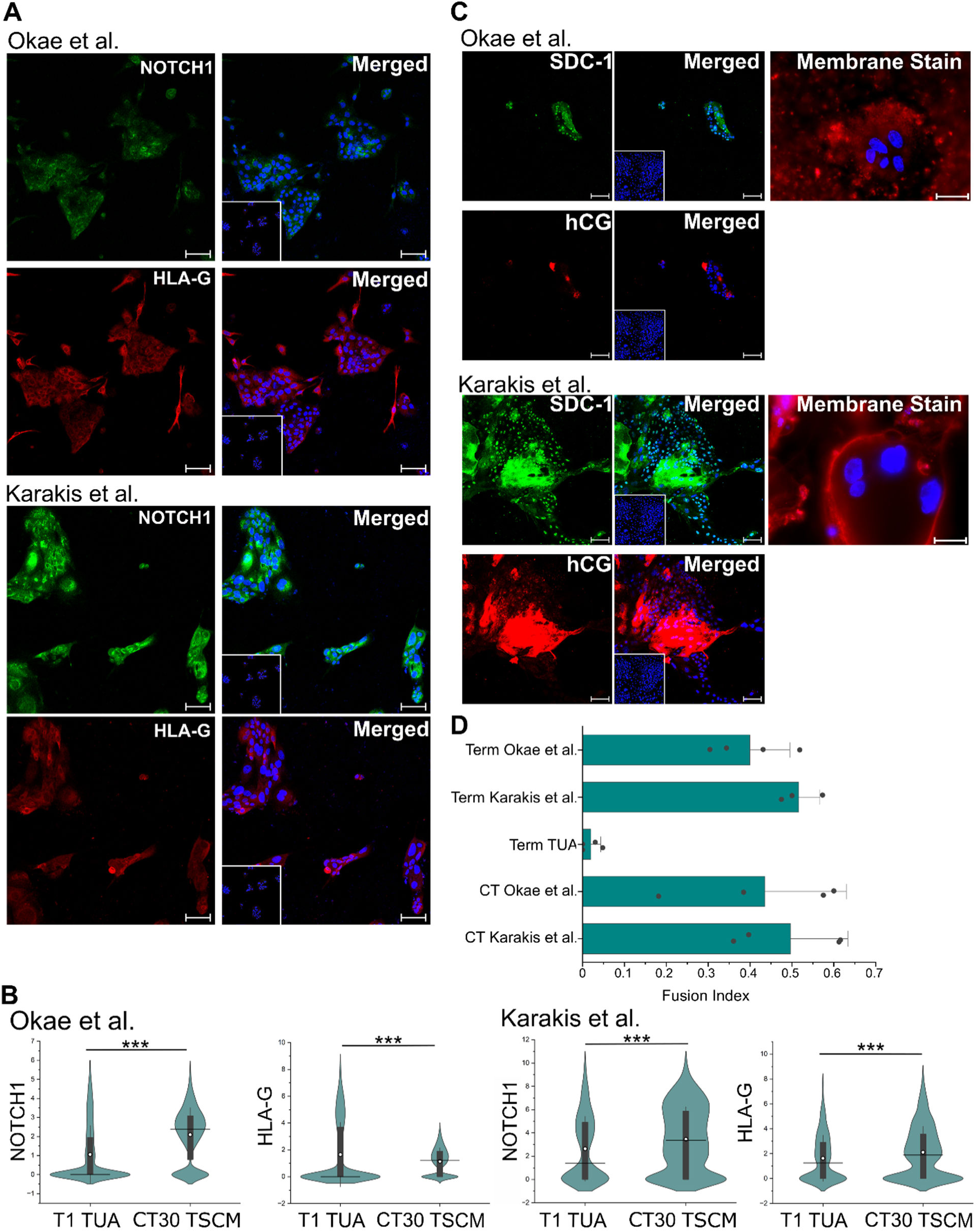
EVT and STB differentiation of hTSCs from term CTBs in TUA medium. (A) Confocal microscopy imaging of T1 hTSCs cultured in TUA and differentiated for 6 days using protocols by Okae et al. or Karakis et al., staining for NOTCH1 and HLA-G at day 6 of differentiation. Nuclei were stained with DAPI. Inset image is isotype control. (B) Quantification of NOTCH1 and HLA-G expression from T1 hTSCs and CT30s differentiated with protocols by Okae et al. or Karakis et al. Data from two biological replicates. White circle represents the mean and the black line represents the median. For T1 hTSCs cultured in TUA using protocol by Okae et al., n= 10992; Karakis et al., n=844. For CT30 hTSCs using protocol by Okae et al., n=5036; Karakis et al., n=1436. (***p-value < 0.001, n.s. = not statistically significant (p>0.05)). Data for EVT differentiation using protocol by Okae et al. have been obtained from our previously published work (15) and re-analyzed for this figure. (C) Di-8-ANEPPS membrane staining and confocal microscopy imaging staining for hCG and SDC-1 at day 6, for T1 hTSCs differentiated to STB using protocols by Okae et al. or Karakis et al. Nuclei were stained with DAPI. Inset images are isotype control. (D) Fusion Index of STB from hTSCs from term (T1 and T2) and first trimester (CT29 and CT30) CTBs, differentiated using protocols by Okae et al. or Karakis et al. T1 and T2 hTSCs in TUA medium are used as a control. The fusion index was calculated as (N-S)/T, where N is the number of nuclei in syncytia, S is the number of syncytia, and T is the total count of nuclei in both fused and unfused cells. Analysis was conducted using at least 7 images, for each biological replicate. All syncytia had a minimum of 3 nuclei within them. Data for STB differentiation using protocol by Okae et al. have been obtained from our previously published work (15) and re-analyzed for this figure. Scale bars are 100 µm for all images.

We have previously described chemically defined conditions for STB differentiation of hTSCs in the absence of forskolin (15). Under these conditions, a membrane stain showed that T1 and T2 hTSCs formed multinucleate cells (**Fig. 2C, S2F**). The fusion indices for the multinucleate cells formed from T1 and T2 hTSCs were comparable to that previously reported for CT29 and CT30 hTSCs under identical differentiation conditions (**Fig. 2D**). Additionally, the differentiated cells expressed the STB markers hCG and SDC-1 (**Fig. 2C, S2E**). Interestingly, considerable cell death was observed when T1 and T2 hTSCs were differentiated in the presence of forskolin, using the protocol described by Okae et al (5). Nevertheless, fusion indices in the surviving cells were comparable to that seen in case of CT29 and CT30 hTSCs differentiated in the presence of forskolin (**Fig. 2D**), and we observed expression of the STB markers hCG and SDC-1 (**Fig. 2C, S2E**).

Taken together, our results show that hTSCs can be derived from term CTBs in TUA medium. These hTSCs can be extensively passaged in TUA medium and retain the ability to differentiate to EVT and STB lineages.

### hTSCs from term placentas can be transitioned into TSCM and cultured extensively and retain differentiation potential

We observed that stable hTSC cultures could not be established from CD49f+ CTBs isolated from term placentas in TSCM, even though cells could be maintained over a few passages. During these early passages, we observed expression of the pan-trophoblast marker KRT7, and markers associated with hTSCs such as GATA3, TEAD4, TFAP2C, YAP, and p63 (**Fig. S4A**). Interestingly, we also observed expression of CDX2; quantitative image analysis comparing expression levels of CDX2 and p63 relative to hTSCs from term CTBs in TUA medium is shown in **Fig. S4B**. Further, we observed that cells in TSCM at early passage numbers did not efficiently differentiate to EVT or STB lineages using protocols described by Okae et al. or chemically defined differentiation conditions described by Karakis et al; expression of the EVT markers Notch1 and HLA-G, or the STB markers SDC-1 and hCG, were low or not observed using either set of differentiation conditions (**Fig. S4C-D**). These results are consistent with the findings reported by Okae et al. that hTSCs could not be derived from term CTBs in TSCM (5).

However, strikingly, we observed that T1 and T2 hTSCs cultured in TUA medium for 5 passages could be transitioned to TSCM and cultured for 20+ passages. In TSCM, T1 and T2 hTSCs retained expression of markers associated with hTSCs, including GATA3, TFAP2C, YAP, TEAD4 and p63 (**Figs. 3A, S5A**). T1 and T2 hTSCs transitioned into TSCM also retained expression of CDX2, although quantitative image analysis showed lower expression of CDX2 and p63 in TSCM relative to cells in TUA medium (**Figs. 3B, S5B**).

**Figure 3:**
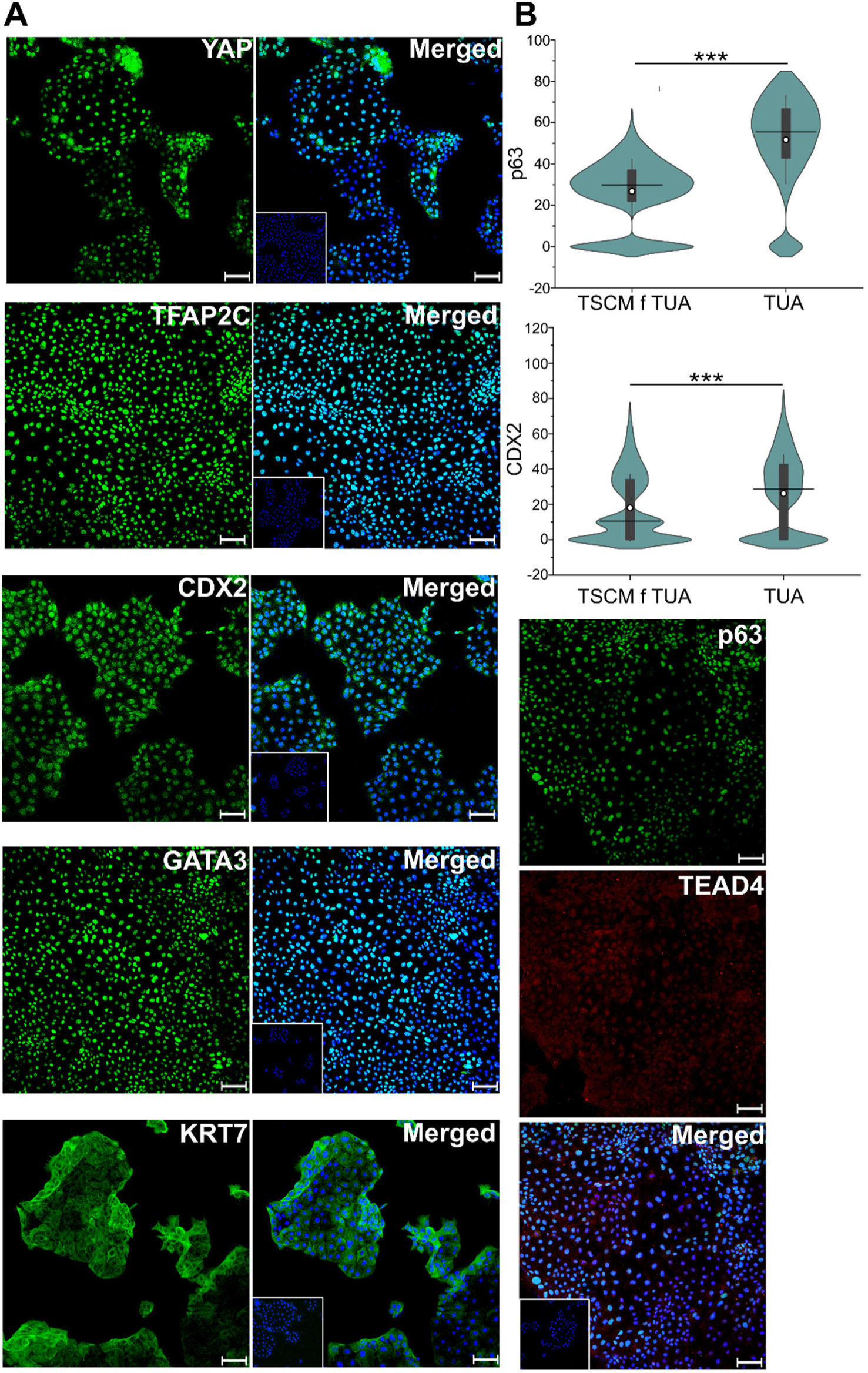
Expression of hTSC markers in cells transitioned to TSCM. (A) Confocal microscopy imaging of T1 hTSCs transitioned to TSCM, staining for TFAP2C, YAP, GATA3, KRT7, p63, TEAD4, and CDX2. p63 was co-stained with TEAD4. Nuclei was stained with DAPI. Inset images are isotype control. Scale bars are 100 µm. (B) Quantification of expression of p63 and CDX2 from T1 hTSCs transitioned into TSCM (n=9931 for p63, n=9210 for CDX2). Data from two biological replicates used; data from T1 hTSCs in TUA medium (same data as in Fig. 1) is shown for comparison. White circle represents the mean and the black line represents the median (***p-value < 0.001). Scale bars are 100 µm for all images.

T1 and T2 hTSCs transitioned into TSCM also retained the ability to differentiate into EVTs and STB. As in the case for cells in TUA medium, we differentiated T1 and T2 hTSCs in TSCM to EVT lineage using protocols described by Okae et al. (5) and Karakis et al (15). Differentiated cells expressing the EVT markers Notch1 and HLA-G were obtained using both protocols (**Figs. 4A, B S6A, B**). Like the case for TUA medium, T1 and T2 hTSCs transitioned to TSCM formed multinucleate cells when differentiated under chemically defined conditions in the absence of forskolin (protocol by Karakis et al.) (**Figs. 4C, S6C**). However, forskolin-mediated STB differentiation of T1 and T2 hTSCs transitioned into TSCM, using the protocol described by Okae et al., resulted in considerable cell death, like the results observed for T1 and T2 hTSCs in TUA medium. Nevertheless, we observed expression of the STB markers hCG and SDC-1 (**Figs. 4C, S6C**), and fusion indices in the surviving cells were comparable to that seen in case of hTSCs in TUA medium differentiated using the same protocol (**Fig. 4C**).

**Figure 4:**
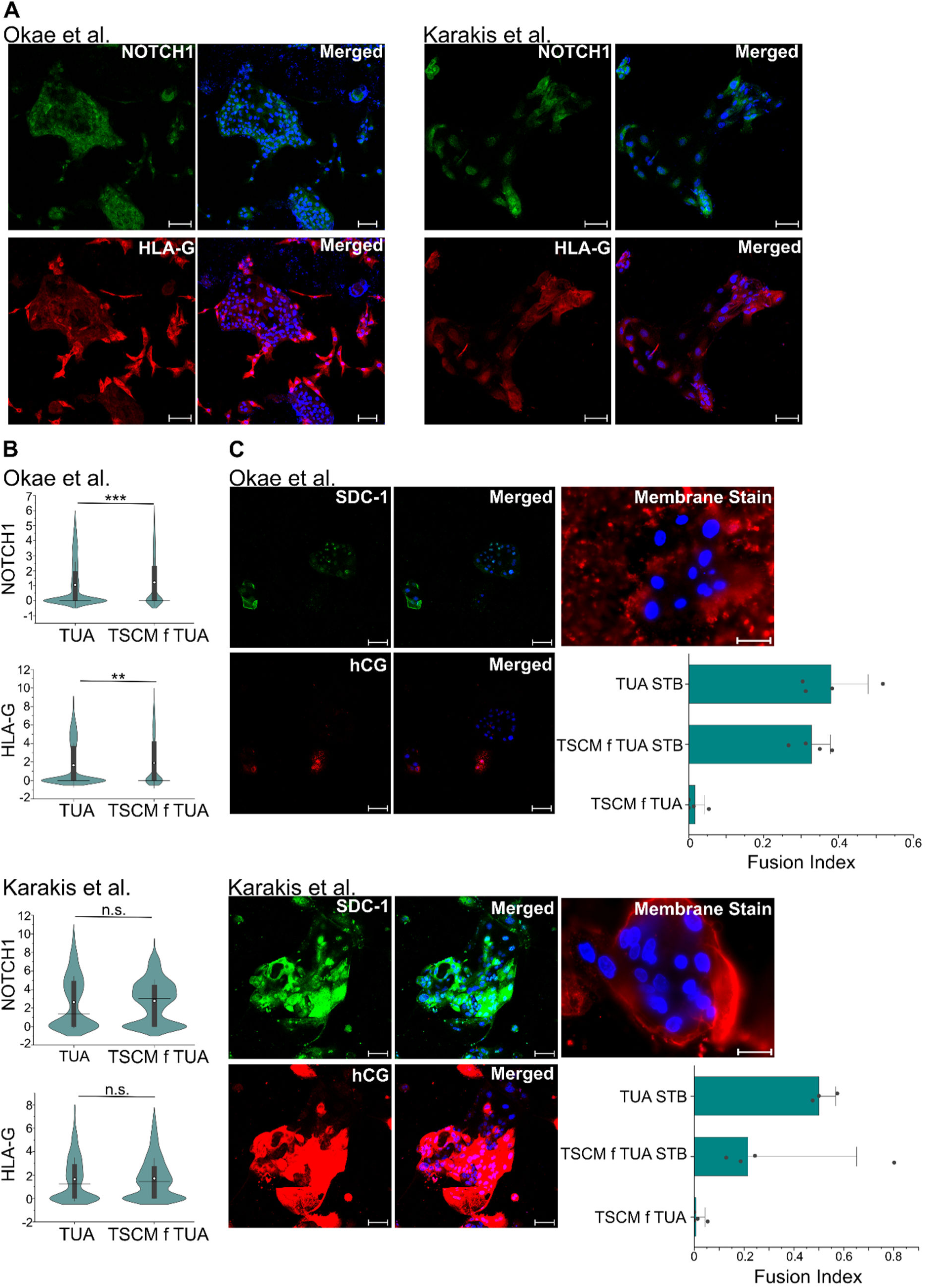
EVT and STB differentiation of hTSCs transitioned into TSCM. (A) Confocal microscopy imaging of T1 hTSCs transitioned into TSCM and differentiated for 6 days using protocols by Okae et al. or Karakis et al., staining for NOTCH1 and HLA-G at day 6 of differentiation. Nuclei were stained with DAPI. (B) Quantification of NOTCH1 and HLA-G expression from T1 hTSCs transitioned into TSCM and differentiated with protocols by Okae et al. (n=2986) or Karakis et al. (n=1080). Data from two biological replicates. Data from T1 hTSCs in TUA medium is shown for comparison (same data as in Fig. 2). White circle represents the mean and the black line represents the median. (**p-value < 0.01, *p-value < 0.05, n.s. = not statistically significant (p>0.05)). (C) Di-8-ANEPPS membrane staining and confocal microscopy imaging staining for hCG and SDC-1 at day 6, for T1 hTSCs in TSCM, differentiated to STB using protocols by Okae et al. or Karakis et al. Nuclei were stained with DAPI. Fusion Index of STB from hTSCs from term placentas (T1 and T2) in TSCM, compared with STB from T1 and T2 hTSCs in TUA medium (same data as Fig. 2), differentiated using protocols by Okae et al. or Karakis et al. T1 and T2 hTSCs in TSCM medium are used as a control. The fusion index was calculated as (N-S)/T, where N is the number of nuclei in syncytia, S is the number of syncytia, and T is the total count of nuclei in both fused and unfused cells. Analysis was conducted using at least 7 images, for each biological replication. All syncytia had a minimum of 3 nuclei within them. Scale bars are 100 µm for all images.

Taken together, these results show that even though hTSCs cannot be derived in TSCM from term CTBs, hTSCs derived in TUA medium can be transitioned into TSCM and cultured extensively thereafter. Further, hTSCs from term CTBs transitioned into TSCM retain their ability to undergo differentiation to EVT and STB lineages.

### Transcriptome analysis of hTSCs derived from term CTBs

We used genome wide RNA-Seq analysis to characterize the transcriptome of T1 and T2 hTSCs cultured in TUA medium or hTSCs transitioned into TSCM from TUA medium (denoted TSCM f TUA). Using data from the literature, we compared the transcriptomes of T1 and T2 hTSCs in these conditions, CT29 and CT30 hTSCs cultured in TSCM (8), hTSCs derived from term CTB in TSCM at 1% oxygen concentration by Wang et al. (14), and primary villous CTBs (vCTBs) isolated from the first trimester (6); CT29 and CT30 hTSCs were used as the control group for comparison in all analysis. A comprehensive list of differentially expressed genes (|log_2_ fold-change| ≥1) is included in **Tables S1-S3**.

Transcript expression for a targeted panel of genes associated with vCTB or early vCTB differentiation; relative expression of mRNA compared to 1st trimester hTSCs. “o” indicates relative expression levels where |log_2_ (fold-change) (FC)| ≥ 1 and FDR p-value < 0.05. Blank grids indicate FDR p-value> 0.05, i.e. not significant

Principal component analysis (PCA) (**Fig. 5A**) and hierarchical clustering analysis (**Fig. 5B**) showed high transcriptome similarity between T1 and T2 hTSCs in TUA or TSCM f TUA and CT29 and CT30 hTSCs, although both have differences relative to primary vCTBs from the first trimester. On the other hand, hierarchical clustering analysis revealed distinct differences between hTSCs derived from term CTBs at 1% oxygen and the other hTSC samples. Notably, 2664 genes are differentially regulated in hTSCs derived at 1% oxygen relative to CT29 and CT30 hTSCs, as compared to 1670 and 1440 genes for T1 and T2 hTSCs in TUA medium and TSCM f TUA respectively (**Fig. 5C**). We further investigated a targeted panel of genes associated with vCTB or vCTB undergoing early differentiation as described(16) (**Fig. 5D**). We did not observe substantial differences in transcript expression of these genes between T1 and T2 hTSCs in TUA and TSCM f TUA relative to CT29 and CT30 hTSCs, with similar expression of the established trophoblast and vCTB markers *GATA2*, *GATA3*, *EGFR*, *YAP1*, *CDH1*, and *TP63* (|log_2_ fold-change| < 1). However, there were some differences observed in expression of markers associated with proliferative trophoblasts (vCTB-p) or markers associated with fusing vCTBs (vCTB-fusing) or cytotrophoblast cell columns (vCTB-ccc); nevertheless, greater differences were observed in hTSCs derived under 1% oxygen.

**Figure 5:**
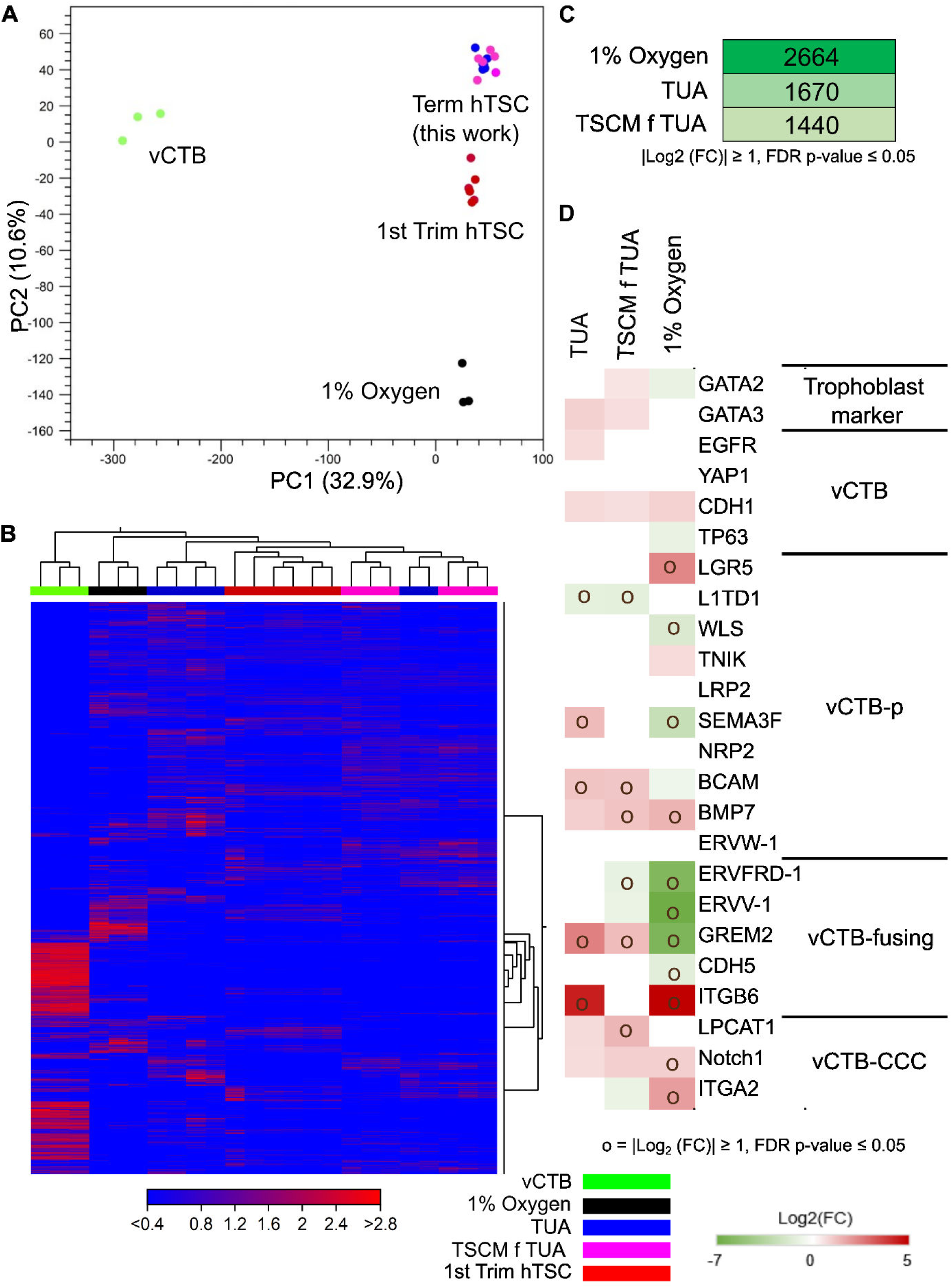
Transcriptome analysis of hTSCs derived from term CTBs. (A) Principal component analysis (PCA) of transcriptome data from primary villous CTB (vCTB), CT29 and CT30 hTSCs cultured in TSCM (1st Trim hTSC), T1 and T2 hTSCs from term CTBs cultured in TUA medium and TSCM f TUA (term hTSC), and hTSCs from term CTB derived in TSCM at 1% Oxygen (1% Oxygen). (B) Hierarchical clustering analysis of transcriptome data for the same samples used in PCA analysis. (C) Number of differentially expressed genes (DEG) for term hTSCs in TUA medium and TSCM f TUA, and hTSCs derived from term CTBs under 1% Oxygen, with false discovery rate (FDR) p-value < 0.05 and |log_2_ (fold-change) (FC)| ≥ 1. 1st trimester hTSCs (CT29, CT30) were used as the control group.

We further used Ingenuity Pathway Analysis (IPA) to assess pathways that are differentially regulated in T1 and T2 hTSCs in TUA, TSCM f TUA, and hTSCs from term CTBs derived under 1 % oxygen, relative to CT29 and CT30 hTSCs (**Fig. 6A-D; Tables S4-S6**). IPA analysis shows that hTSCs derived in 1% oxygen exhibit significant upregulation of DNA damage-induced and oxidation stress-induced senescence pathways (**Fig. 6A**). Further, Wnt and the pre-NOTCH expression and processing pathway, which activates NOTCH signaling downstream, were upregulated in these cells relative to CT29 and CT30 hTSCs. Notably NOTCH signaling is a key determinant of differentiation to the EVT lineage and markers for cytotrophoblast cell columns (vCTB-CCC) are upregulated in hTSCs derived in 1% oxygen (**Fig. 5D**). Interestingly, these pathways were not upregulated in T1 and T2 hTSCs (**Fig. 6B, C**); rather, we observed upregulation of interferon alpha/beta signaling in hTSCs in both TUA medium and TSCM f TUA. Additionally, interferon, interferon gamma, and ISGylation signaling pathways were upregulated in hTSCs in TUA medium but not in TSCM f TUA. Notably, the ISGylation pathway is associated with lipid metabolism (17–20). Use of lipid-rich albumin is critical to deriving hTSCs from term CTBs, as discussed in the following section, and these lipids are withdrawn when hTSCs are transitioned to TSCM from TUA medium.

**Figure 6:**
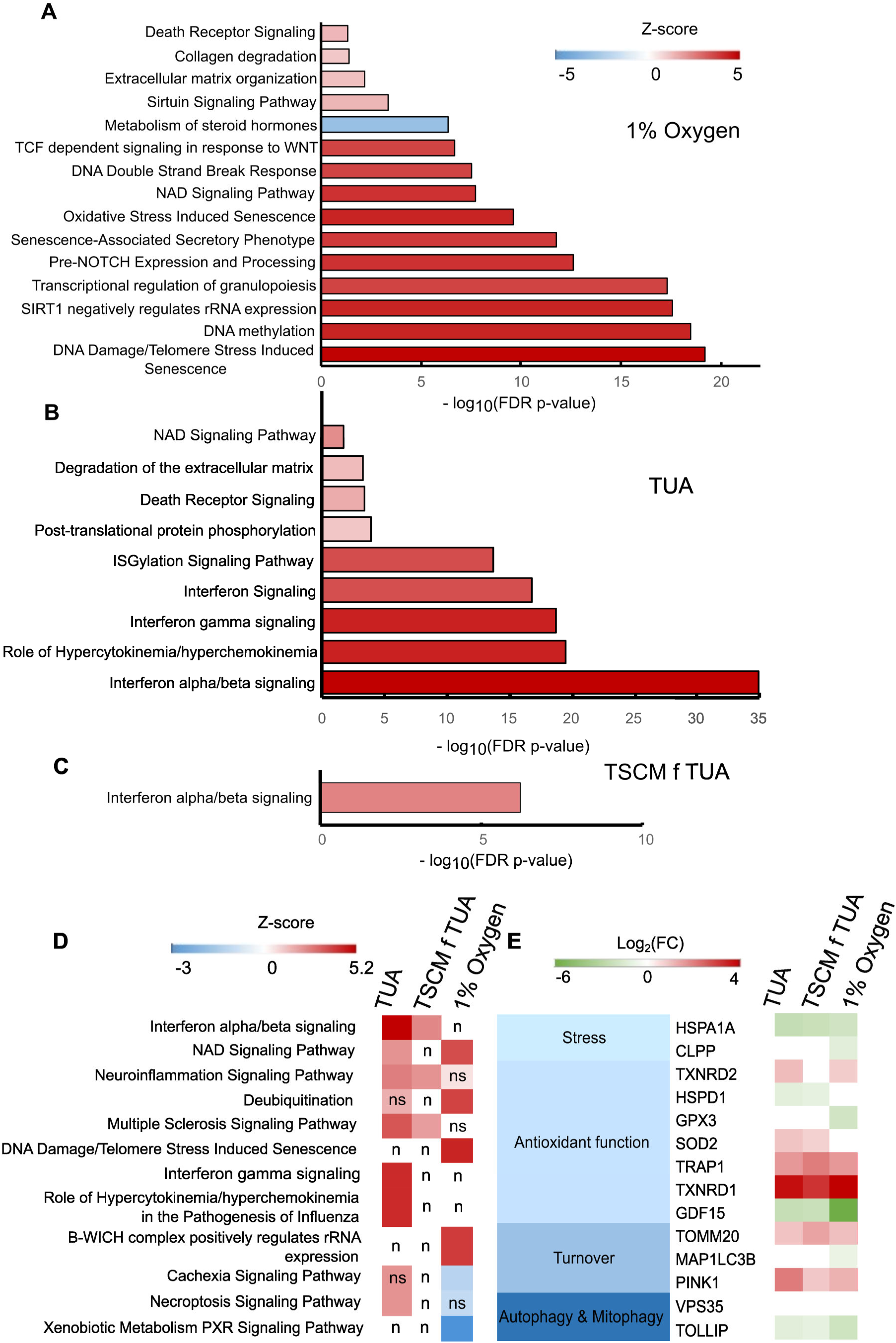
Ingenuity pathway analysis (IPA) of most abundant genes in term hTSCs. (A) – (C) IPA pathway analysis for hTSCs from term CTBs derived under 1% Oxygen (A), and T1 and T2 hTSCs in TUA medium (B) and TSCM f TUA (C). First trimester hTSCs (CT29, CT30) were used as the control group. IPA analysis includes the top five canonical pathways and other significant trophoblast related cellular pathways. Most abundant genes were selected following the cut-off FDR p-value ≤0.05, |log2 (fold-change (FC))| ≥ 2, maximum group mean ≥ 5. (D) Comparative IPA pathway analysis between hTSCs from term CTBs. First trimester hTSCs (CT29, CT30) were used as the control group. IPA analysis includes the top canonical pathways. Most abundant genes were selected following the cut-off FDR p-value ≤ 0.05, |log2 (fold-change (FC))| ≥ 2, maximum group mean ≥ 5. Here, n indicates Z-score is not available for that comparison, ns indicates FDR p-value > 0.05, i.e. non-significant Targeted panel of mitochondrial genes showing differences in relative mRNA expression between hTSCs from term CTBs and hTSCs from first trimester CTBs, with FDR p-value ≤0.05. Blank grids indicate FDR p-value > 0.05, i.e. non-significant

To provide a comprehensive comparison of upregulated and downregulated pathways, we conducted a comparative analysis of IPA outputs from **Fig. 6A-C** across term CTB derived hTSCs (**Fig. 6D**). Additionally, we performed cluster analysis on the differentially expressed genes (DEGs) identified in **Fig. 5B** using the elbow method to define clusters based on their expression patterns under different derivation conditions. These clusters were then subjected to gene ontology (GO) analysis to identify enriched biological pathways. Performing GO analysis on these individual clusters allowed for more specific enrichment results; these are presented in **Tables S7-S19.**

Finally, as discussed in the following section, use of the mitochondrial pyruvate uptake inhibitor UK5099 was essential for derivation of functional hTSCs from term CTBs. Therefore, we investigated a targeted panel of genes associated with mitochondrial regulation (**Fig. 6E**). Specifically, we focused on a set of genes that have been reported to be differentially expressed in primary chorionic tissue in the third trimester relative to the first trimester (21). This previous study reported that expression of the antioxidant genes *CLPP*, *TXNRD2*, *HSPD1* and *GPX3* was lower in tissue from the third trimester relative to the first trimester; on the other hand, *SOD2*, *HSPA1A*, *TRAP1*, *TXNRD1* and *GDF15* were upregulated in tissue from the third trimester. Additionally, expression of the mitophagy and autophagy-related genes *TOMM20*, *MAP1LC3B* and *PINK1* was upregulated in tissue from the third trimester, and *VPS35* and *TOLLIP* expression was downregulated. Our analysis showed that differential expression of these genes was fairly consistent across hTSCs derived from third trimester CTBs (T1 and T2 hTSCs in TUA medium and TSCM f TUA, and hTSCs derived under 1 % oxygen), relative to CT29 and CT30 hTSCs derived from first trimester placentas. However, trends observed in third vs. first trimester primary tissue were not entirely conserved. For instance, similar to primary tissue, *HSPD1* and *TOLLIP* were downregulated and *SOD2*, *TRAP1*, *TXRND1*, *TOMM20* and *PINK1* were upregulated in T1 and T2 hTSCs relative to CT29 and CT30 hTSCs. On the other hand, unlike primary tissue from third vs. first trimester, T1 and T2 hTSCs showed downregulation of *HSPA1A* and *GDF15*; there were no statistically significant differences in expression of *CLPP*, *GPX3*, *MAP1LC3B* and *VPS35* between T1 and T2 hTSCs from term CTBs and CT29 and CT30 hTSCs.

Overall, our transcriptome analysis shows that T1 and T2 hTSCs derived from term CTBs, in both TUA medium and TSCM f TUA, show high transcriptome similarity to CT29 and CT30 hTSCs derived from first trimester placentas. Nevertheless, there are some differences in transcript expression, albeit fewer than those observed for hTSCs derived from term CTBs under 1% oxygen. Collectively with data on extended culture of T1 and T2 hTSCs in TUA and TSCM f TUA (**Figs.1, S1, 3, S5**), and differentiation to EVT and STB lineages (**Figs. 2, S2, 4, S6**), our transcriptome analysis confirms that hTSCs can indeed be derived from term CTBs under atmospheric oxygen conditions using TUA medium. These hTSCs can be cultured extensively in TUA medium and subsequently in TSCM, while retaining differentiation potential.

### Comparative transcriptome analysis of hTSCs derived from term CTBs and hTSCs from human pluripotent stem cells

Several studies have investigated the derivation of hTSCs from human pluripotent stem cells (hPSCs) through reprogramming or differentiation. For instance, Dong et al. (9) derived hTSCs from naïve hPSCs and iPSCs (in 5i/L/A or PXGL media) by transitioning these naïve cells into TSCM. Guo et al. (22) generated hTSCs from hPSCs by inducing trophectoderm differentiation using PD0325901 (MEK inhibitor) and A83-01 (TGF-β/ALK4/5/7 inhibitor) and subsequently culturing the cells in TSCM. Io et al. (23) developed a protocol for naive cytotrophoblast-like cells (nCTs) from naive hPSCs via an intermediate naive trophectoderm-like state (nTEs), followed by expansion of hTSCs in ACE medium. Zorzan et al. (24) established chemically reset TSCs (ccTSCs) from primed iPSCs through a resetting phase by culture in MEKi/LIF/HDACi followed by PXGL medium, before transitioning to TSCM. Finally, in previous work we have shown that treatment of hPSCs with BMP4, S1P or S1PR3 agonist, and SB431542 was sufficient to generate CTB like cells which could be subsequently transitioned to TSCM or TM4 medium to maintain two distinct hTSC populations (8). We compared hTSCs in TUA medium derived from term placentas (this study) with hTSCs derived in these aforementioned studies. **Fig.7** shows a comparative transcriptome analysis using publicly available datasets from these studies (8, 9, 22–24), including PCA analysis (**Fig. 7A**), heatmap analysis (**Fig. 7B**), and trophoblast-specific differentially expressed genes (**Fig. 7C**). For this analysis, we included a subset of hTSC cell lines generated in these studies - hTSCs derived from naïve H9 and WIBR3 hPSCs from Dong et al. (GSE138762) (9); hTSCs derived from hPSC lines HNES1, niPSC2, and niPSC4 lines by Guo et al. (GSE167089) (22); H9 hESC-derived nCTs that go through an intermediate nTE state from Io et al. (GSE144994) (23); ccTSC.02 derived from chemically reset primed iPSCs from Zorzan et al. (GSE184562) (24); and hTSCs in TSCM medium obtained from H1 and H9 hESCs after BMP4 treatment, from Mischler et al. (GSE137295) (8). Unless otherwise specified, first-trimester hTSCs, CT 29 and CT 30 were used as the control group for analysis.

**Figure 7:**
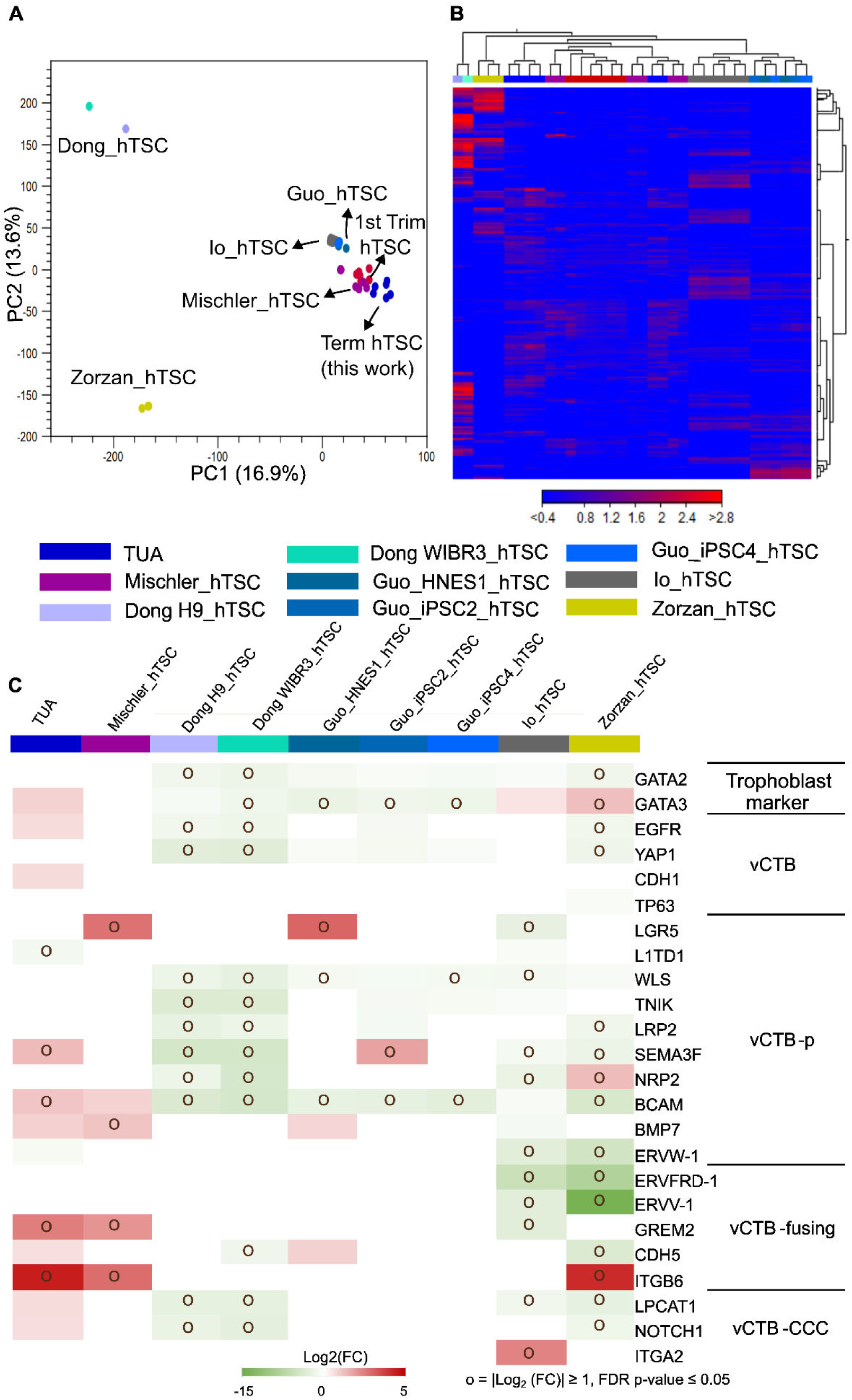
Comparison of transcriptome analysis of hTSCs derived from term CTBs and hTSCs derived from hPSCs. (A) Principal component analysis (PCA) of transcriptome data from T1 and T2 hTSCs from term CTBs cultured in TUA medium, CT29 and CT30 hTSCs cultured in TSCM (1st Trim hTSC), and hTSCs derived from hPSCs. (B) Hierarchical clustering analysis of transcriptome data used in PCA analysis. (C) Transcript expression for a targeted panel of genes associated with vCTB or early vCTB differentiation; relative expression of mRNA compared to 1st trimester hTSCs. “o” indicates relative expression levels where |log2 (fold-change) (FC)| ≥ 1 and FDR p-value ≤ 0.05. Blank grids indicate FDR p-value> 0.05, i.e. not significant. vCTB-p: proliferative vCTB, vCTB-CCC: vCTB – cytotrophoblast cell column

PCA analysis (**Fig 7A**) reveals that term hTSCs in TUA medium cluster closely with hTSCs derived in studies by Guo et al. (22), Io et al. (23), Mischler et al. (8) and first trimester hTSCs derived by Okae et al. (5) In contrast, hTSCs derived by Dong et al. and Zorzan et al. form distinct clusters, suggesting transcriptomic differences between these cells and term hTSCs in TUA medium. A similar clustering pattern is observed in the heatmap analysis in **Fig. 7B**. Further, differential gene expression analysis (**Fig. 7C**) shows the expression patterns of key trophoblast-specific markers of vCTB or trophoblast subtypes representative of early vCTB differentiation as described (16), relative to first trimester hTSCs. There is reasonable similarity between term hTSCs in TUA medium and hTSCs derived from hPSCs. A comprehensive list of differentially expressed genes (|log_2_ fold-change| ≥1) from this analysis is included in **Tables S20-S27**.

Collectively, these analyses show that term hTSCs in TUA medium (and term hTSCs in TSCM f TUA) exhibit transcriptional similarity with both first trimester hTSCs and hTSCs derived from hPSCs.

### Lipid-rich albumin and UK5099 are essential for derivation of hTSCs from term CTBs

We observed that primary CD49f+ CTBs could not be maintained in culture in TSCM supplemented with UK5099 (without AlbuMAX II) during early passages, suggesting that the presence of lipid-rich albumin is essential for establishing hTSC cultures from term CTBs. We further investigated if hTSCs can be derived from term CTBs in TSCM supplemented with AlbuMAX II alone (no UK5099); note that the same primary cells used to derive T1 and T2 hTSCs in TUA medium were used in these studies. We observed that cells could be maintained for several passages under these conditions (TA medium), and expressed relevant CTB markers including, GATA3, P63, CDX2, TFAP2C, KRT7, and YAP (**Figs. 8A, S7A**), like hTSCs cultured in TUA medium. However, cells in TA medium were unable to efficiently differentiate to EVT and STB using either the chemically defined conditions described by Karakis et al. (15) or protocols described by Okae et al (5). HLA-G+ EVT cells were not obtained using either protocol in T1 hTSCs; some HLA-G+ cells were observed in T2 hTSCs when the protocol by Okae et al. was used (**Figs. 8B, C, S7B, C**). Similarly, we did not observe significant and consistent expression of the STB markers hCG and SDC-1 (**Fig. 8D, S7D**). These results show that although hTSC-like cells derived from term CTBs can be maintained in TA medium, they are deficient in their ability to differentiate to EVT and STB.

**Figure 8:**
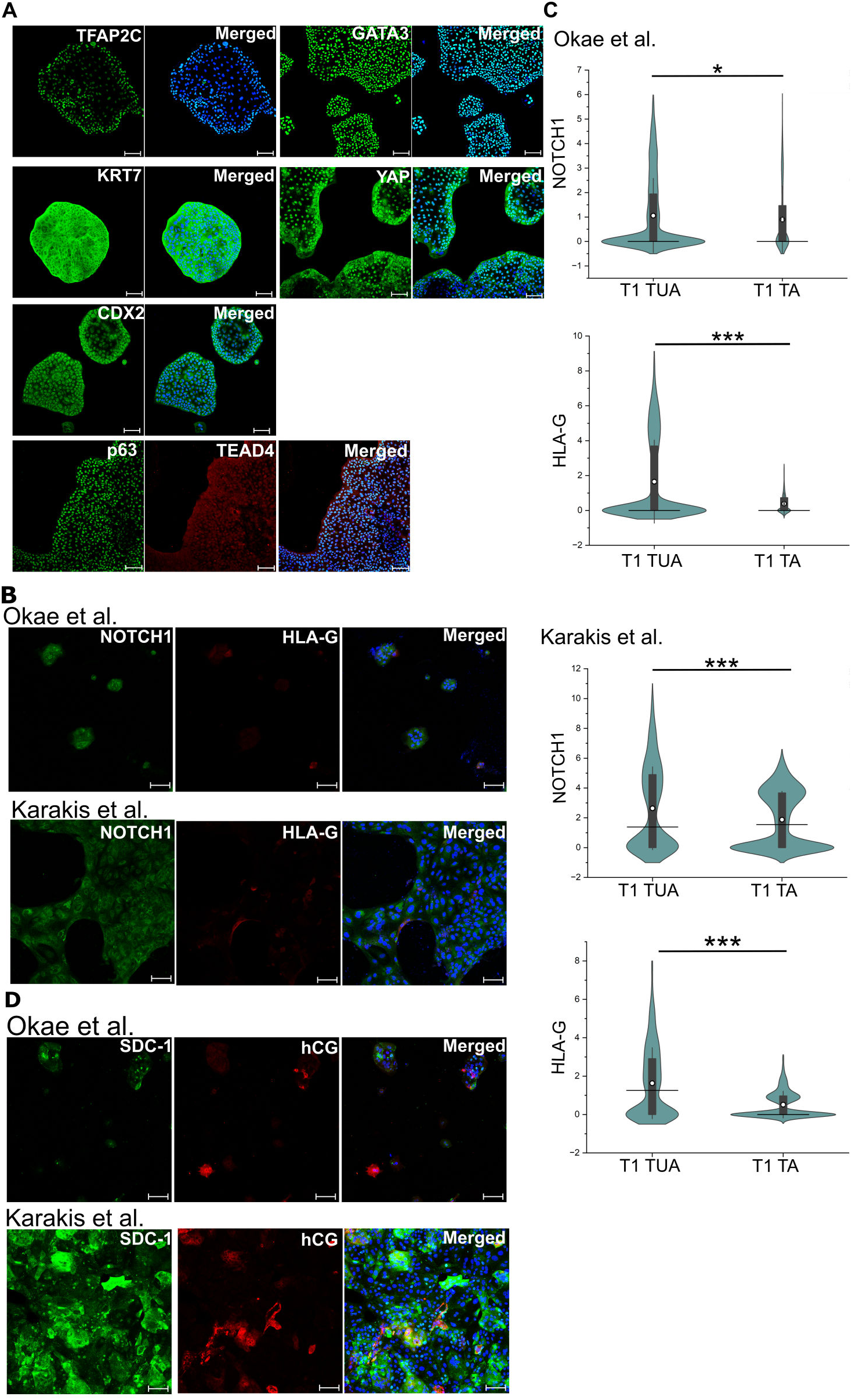
Attempted derivation of hTSCs from term CTBs in TA medium. (A) Confocal microscopy imaging of cells obtained during attempted derivation of hTSCs in TA medium, staining for TFAP2C, YAP, GATA3, KRT7, p63, TEAD4, and CDX2; p63 was co-stained with TEAD4. Nuclei were stained with DAPI. Primary CTBs are from the same placenta as those used for deriving T1 hTSCs in TUA medium. (B) Confocal microscopy imaging of cells obtained during attempted derivation of hTSCs in TA medium, differentiated to EVTs for 6 days using protocols by Okae et al. or Karakis et al., staining for NOTCH1 and HLA-G. Nuclei were stained with DAPI (blue). (C) Quantification of NOTCH1 and HLA-G expression in cells obtained during attempted derivation of term CTBs in TA medium (labeled T1 TA), differentiated with protocols by Okae et al. (n=1356) or Karakis et al. (n=1688). Data from two biological replicates used. Data from T1 hTSCs in TUA medium is shown for comparison (same data as in Fig. 2). White circle represents the mean and the black line represents the median. (***p-value < 0.001, n.s. = not statistically significant (p>0.05)). (D) Confocal microscopy imaging of cells obtained during attempted derivation of hTSCs in TA medium, differentiated to STB for 6 days using protocols by Okae et al. or Karakis et al., staining for SDC-1 and hCG. Nuclei were stained with DAPI (blue).

To further examine the role of UK5099 in hTSC derivation from term CTBs, we used RNA-Seq analysis to compare the transcriptome of hTSC-like cells in TA medium and T1 and T2 hTSCs in TUA medium. The comprehensive list of differentially regulated genes is included in **Table S28**. In particular, we focused on the targeted panel of genes associated with the vCTB or vCTB early differentiation as described (16) (**Fig. 9A**). We observed that genes associated with proliferative vCTB are decreased in hTSC-like cells in TA medium. Nevertheless, we see high transcriptome similarity between cells in TA and TUA medium. We also used Ingenuity Pathway Analysis to identify signaling pathways that are differentially regulated in hTSC-like cells in TA medium relative to hTSCs in TUA medium (**Table S29**). Interestingly, removal of the mitochondrial pyruvate uptake inhibitor UK5099 from TUA medium resulted in downregulation of ISGylation, interferon, and interferon alpha/beta signaling pathways (**Fig. 9B**), suggesting a role for mitochondrial metabolism in upregulation of these pathways in TUA medium.

**Figure 9:**
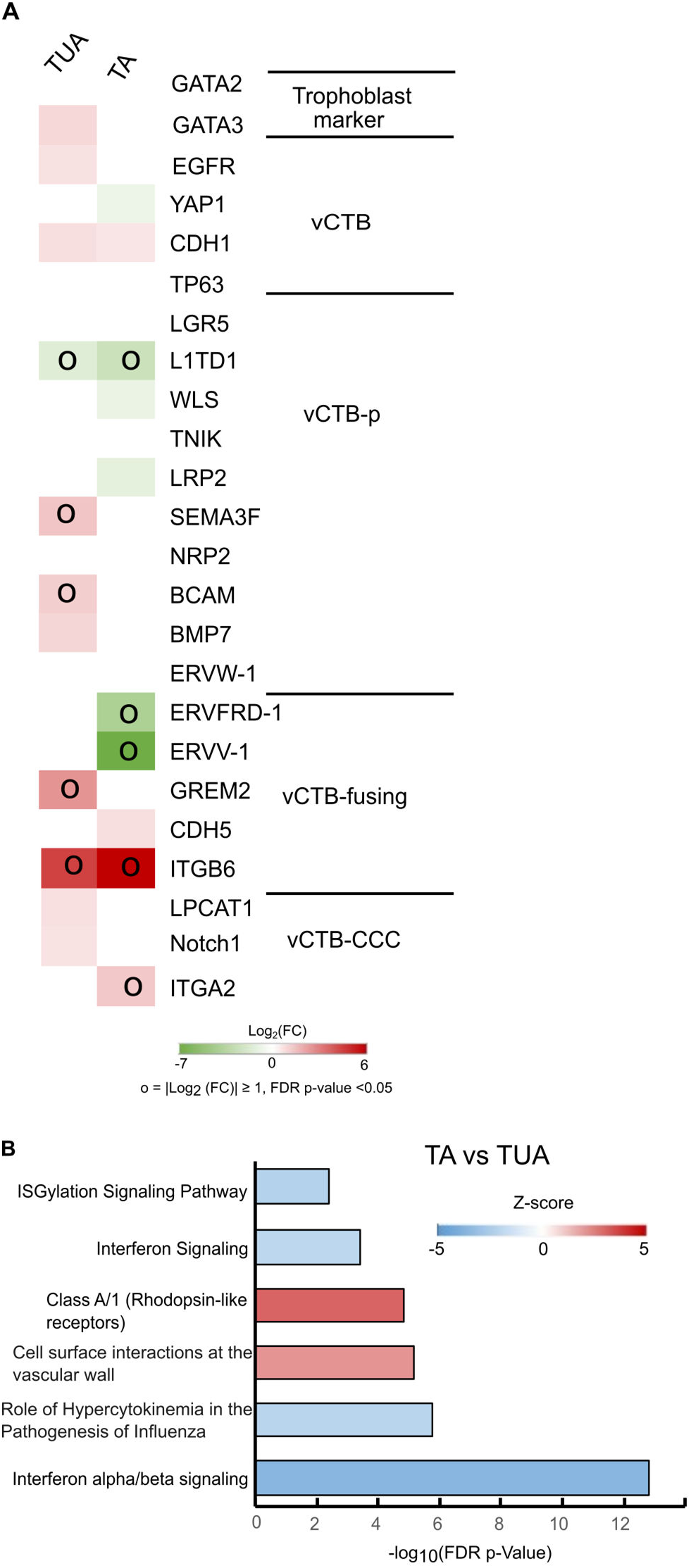
Transcriptome analysis of hTSCs derived from term CTBs in TA. (A) Transcript expression for a targeted panel of genes associated with vCTB or early vCTB differentiation in T1 and T2 hTSCs in TUA medium or cells obtained during attempted derivation of hTSCs in TA medium; relative expression of mRNA compared to 1st-trimester hTSCs (CT29, CT30). o indicates relative expression levels where |log2 (fold-change (FC))| ≥ 1 and FDR p-value < 0.05. Blank grids indicate FDR p-value> 0.05, i.e. non-significant. (B) IPA pathway analysis for cells obtained during attempted derivation of hTSCs in TA medium; primary CTBs are from the same placentas used for derivation of T1 and T2 hTSCs. T1 and T2 hTSCs cultured in TUA medium were used as the control group. IPA analysis includes the top five canonical pathways and other significant trophoblast-related cellular pathways. Most abundant genes were selected for IPA analysis with a more stringent dataset following the cut-off FDR p-value ≤0.05, |log2(fold-change (FC))| ≥ 2, maximum group mean ≥ 5.

Previous studies have reported that lipid-rich albumin contains phospholipids such as lysophosphatidic acid (LPA) and free fatty acids (25). We further investigated if AlbuMAX II could be replaced by LPA or chemically defined lipid concentrate (CDL). Accordingly, we tried to derive hTSCs in TSCM supplemented with UK5099 and LPA (TUL medium) or CDL (TUC medium). Culture of cells in these media was far less robust than in TUA medium, limiting their analysis; indeed, expansion of cells frozen after culture of cells in TUC medium for few passages was difficult. Nevertheless, our analysis showed that cells expressing the hTSC markers GATA3 and CDX2 could be cultured in TUL medium, although they expressed lower levels of TEAD4 and exhibited greater non-nuclear staining of p63 and YAP (**Fig. S8A, B**). Similarly, cells expressed CDX2 in TUC medium but showed low expression of GATA3, TEAD4, TFAP2C, and p63 (**Fig. S8C, D**). However, cells in both TUL and TUC medium showed deficient differentiation to EVT and STB using two different differentiation protocols (**Fig. S9**); we did not consistently observe cells with the EVT markers HLA-G and Notch1 or the STB markers hCG and SDC-1. Thus, functional hTSCs could not be derived when the lipid-rich albumin in TUA medium was replaced by LPA or CDL.

Taken together, our results show that lipid-rich albumin is critical for culture and expansion of term CTBs. However, addition of the mitochondrial pyruvate uptake inhibitor UK5099 is necessary for deriving hTSCs that retain differentiation potential.

## Discussion

In this study, we have shown that hTSCs can be derived from term CTBs under atmospheric oxygen conditions in TUA medium; TUA medium is TSCM used for derivation of hTSCs from first trimester placentas and blastocyst-stage embryos described by Okae et al., supplemented with lipid-rich albumin (AlbuMAX II) and a low concentration (50 nM) of the mitochondrial pyruvate uptake inhibitor UK5099. We derived two hTSC lines, T1 (female) and T2 (male) from CD49f+ CTBs isolated from term placentas with no known pathology. These hTSCs in TUA medium express markers associated with undifferentiated hTSCs and can differentiate to EVT and STB. Strikingly, hTSCs from term CTBs cultured in TUA medium for a few passages can be transitioned into TSCM and extensively cultured further. Importantly, hTSCs transitioned from TUA medium into TSCM retain their ability to differentiate to EVT and STB. Finally, hTSCs from term CTBs in TUA medium as well those transitioned into TSCM exhibit high transcriptome similarity to hTSCs derived from first trimester CTBs.

### Role of lipid-rich albumin

TSCM contains 0.2% fetal bovine serum and 0.3% fatty acid free bovine serum albumin (BSA). Although term CTBs could be cultured for a few passages in TSCM under atmospheric oxygen conditions, they could not be maintained in long-term culture. However, cells could be extensively cultured in TSCM supplemented with lipid-rich albumin (TA medium), although these cells were deficient in differentiation potential. Strikingly, Yang et al. (13) modified the medium described by Turco et al. (7) by supplementation with 10% FBS for derivation of trophoblast organoids from term placentas; FBS contains lipid-rich albumin. Collectively, these results suggest that lipid supplementation plays an important role in enabling derivation of self-renewing cultures (hTSCs or organoids) from late gestation placentas. In this context, Wang et al. have derived hTSCs from term placentas in TSCM – without additional lipid supplementation – at 1% oxygen (14); notably, hypoxia has been reported to upregulate lipid synthesis (26). Yet, it is important to note that lipid supplementation can be withdrawn after a few passages of hTSC derivation in TUA medium; hTSCs can subsequently be cultured in TSCM. Therefore, we speculate that lipid-rich albumin is necessary for initial adaptation of term CTBs – but not CTBs from early gestation – in culture, and to overcome stress associated with transfer of primary cells to an in vitro cell culture environment. Consistent with this speculation, lipid supplementation has been shown to increase the efficiency of primary and secondary intestinal organoid formation from old mice (27).

### Role of mitochondrial pyruvate uptake inhibitor

Although hTSC-like cells can be derived from term CTBs in TSCM supplemented with lipid-rich albumin, these cells are deficient in their ability to differentiate to EVT and STB. Thus, supplementation with the mitochondrial pyruvate uptake inhibitor UK5099 is critical for derivation of hTSCs from term CTBs in our conditions. We have used a low concentration of UK5099 in our study (50 nM; IC50 is 50 nM); we observe significant cell death when high concentrations of UK5099 (greater than 10x IC_50_) are used with hTSCs in TSCM. Notably, at concentrations similar to that used here, UK5099 has been shown to increase mitochondrial respiration through lipid cycling in brown adipocytes (28). Our transcriptome analysis also shows that interferon-related signaling pathways, including ISGylation, are upregulated in TUA medium relative to TA medium. Pertinently, reduction in levels of ISG15 that is required for ISGylation is associated with preeclampsia and knockdown of ISG15 expression impairs invasion of HTR8/SVneo cells in culture (29). ISGylation has also been reported to modulate the stability of multiple proteins, including YAP, P63, and HIF1α, which play a critical role in trophoblast development (30). ISGylation is also important to maintain efficient oxidative phosphorylation in pancreatic cancer stem cells (19). Further studies are needed to identify the specific molecular mechanisms through which UK5099 exerts its effects on hTSCs in TUA medium.

### Comparison of hTSCs from term placentas with hTSCs from early gestation

Our analysis shows high transcriptome similarity between hTSCs from term CTBs derived in TUA medium and hTSCs (CT29, CT30) from early gestation. Further, hTSCs in TUA medium can be transitioned into the same medium used for culture of hTSCs from early gestation. Like hTSCs in TUA medium, the transitioned hTSCs in TSCM exhibit high transcriptome similarity to early gestation hTSCs and retain the ability to differentiate into EVT and STB. Moreover, T1 and T2 hTSCs derived in this study showed protein expression of the early trophoblast marker CDX2, unlike CT29 and CT30 hTSCs although upregulation of *CDX2* transcript was not found to be statistically significant in RNA-Seq analysis (log_2_ fold-change = 1.29 for comparison between T1 and T2 hTSCs vs CT29 and CT30 hTSCs, FDR p-value = 0.14). These results are consistent with observations by Wang et al. who report high transcriptome similarity between hTSCs derived at 1% oxygen and hTSCs from early gestation (14). Similarly, Yang et al. show that the transcriptome of trophoblast organoids derived from term placentas is similar to first trimester placental tissue (13). Strikingly, we also observe transcriptome similarity between T1 and T2 hTSCs and hTSCs derived from hPSCs in several different studies (**Fig. 7**). Collectively, these studies raise the intriguing possibility that derivation of hTSCs from term CTB results in reprogramming to an earlier developmental state. Such reprogramming is likely dependent on epigenetic modifications upon culture of term CTBs. Indeed, valproic acid, a known epigenetic modifier, is a key component of TSCM. Further studies are needed to rigorously assess epigenetic differences between hTSCs derived from term placentas, hTSCs from early gestation, and primary CTBs across gestation.

In conclusion, we have shown that functional hTSCs can be derived in a straightforward manner from placentas at birth using TUA medium, i.e. TSCM supplemented with UK5099 and AlbuMAX II. Importantly, our approach does not require equipment for control of oxygen concentrations and once derived, hTSCs can be transitioned into TSCM that is widely used for culture of hTSCs from first trimester placentas and blastocyst stage embryos. These results will enable facile derivation of hTSCs from normal and pathological placentas at birth and enable mechanistic studies that address critical knowledge gaps in our understanding of human placental development.

## Methods

### Key resources

**Table 1.**
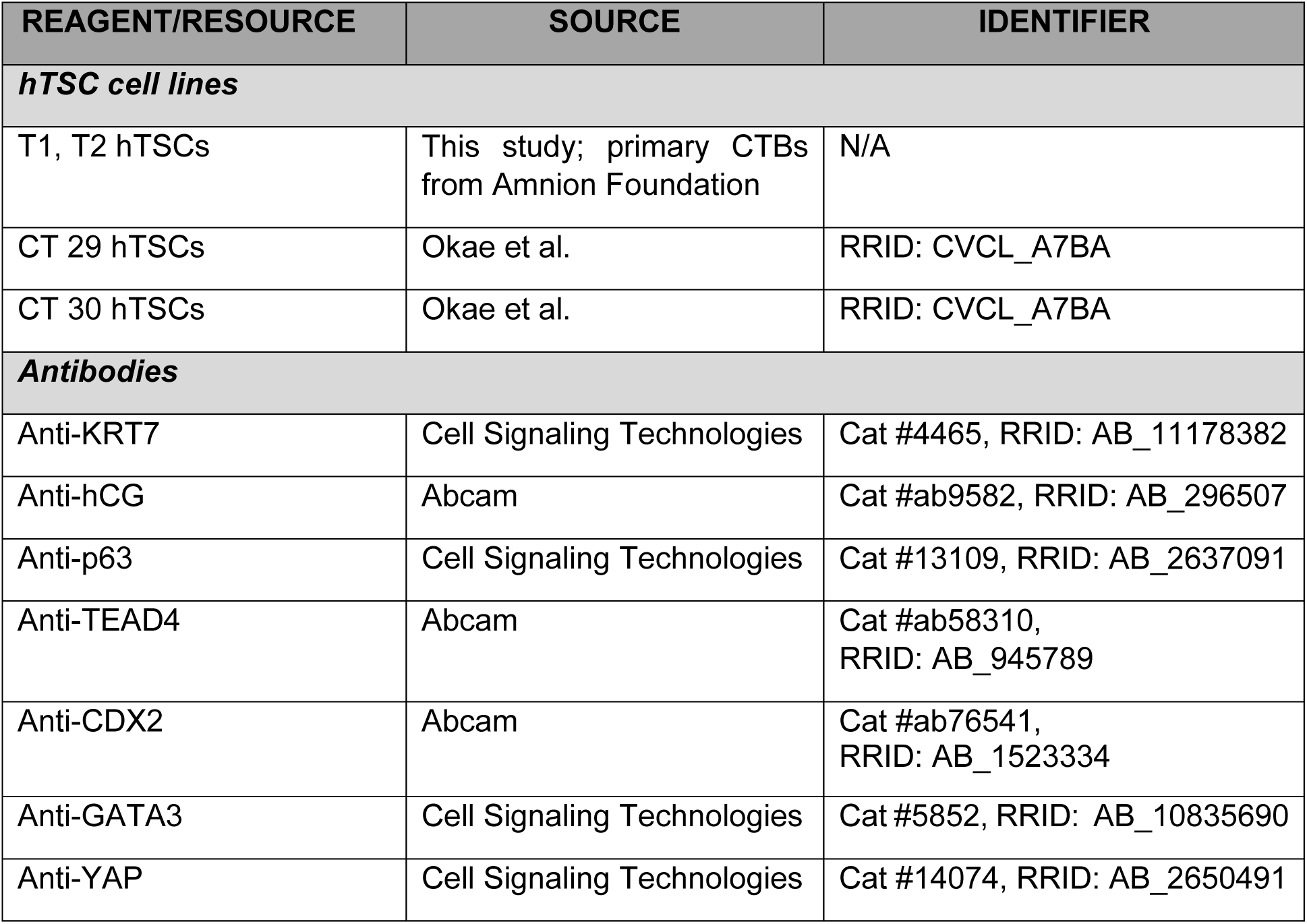
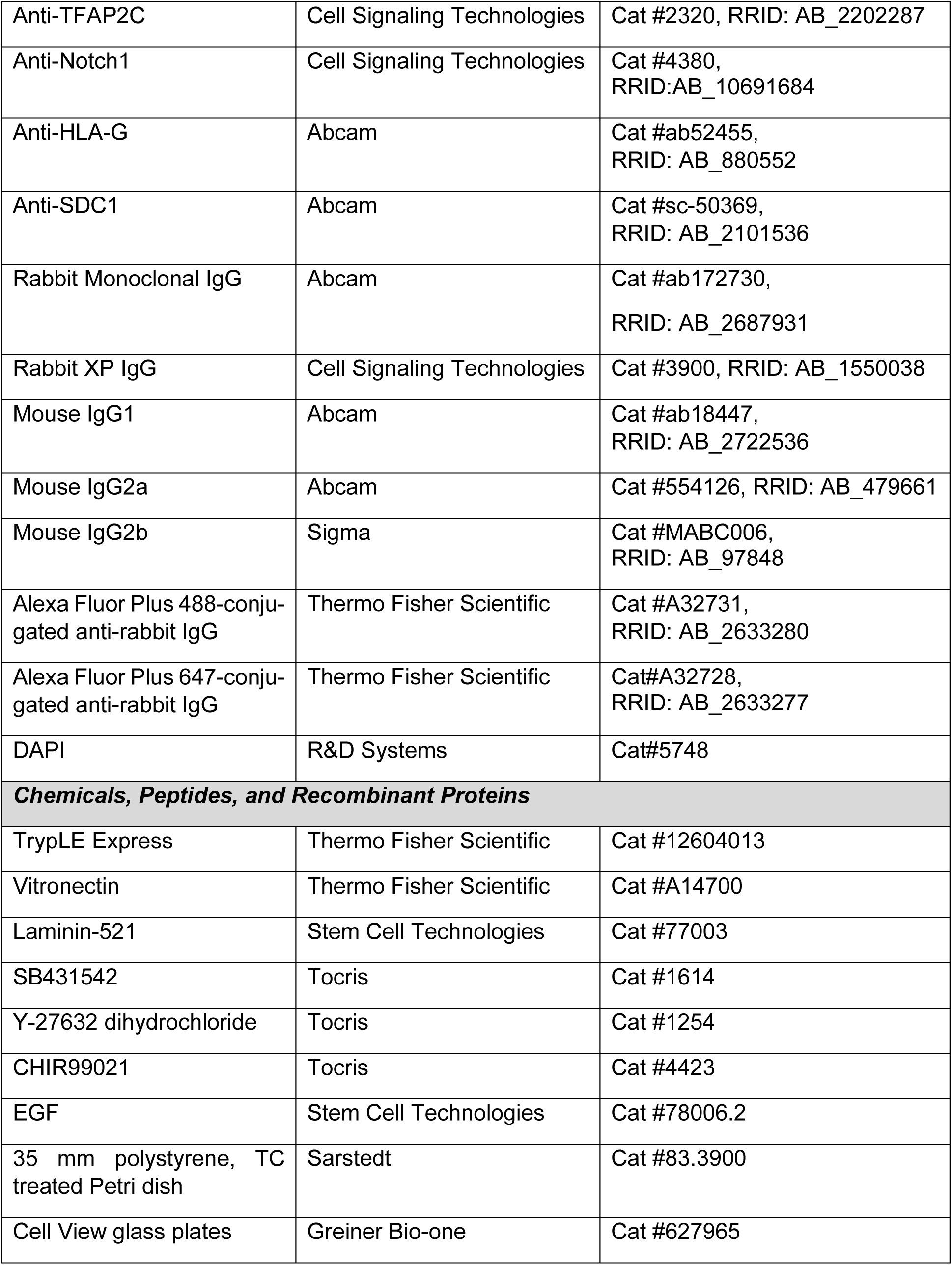
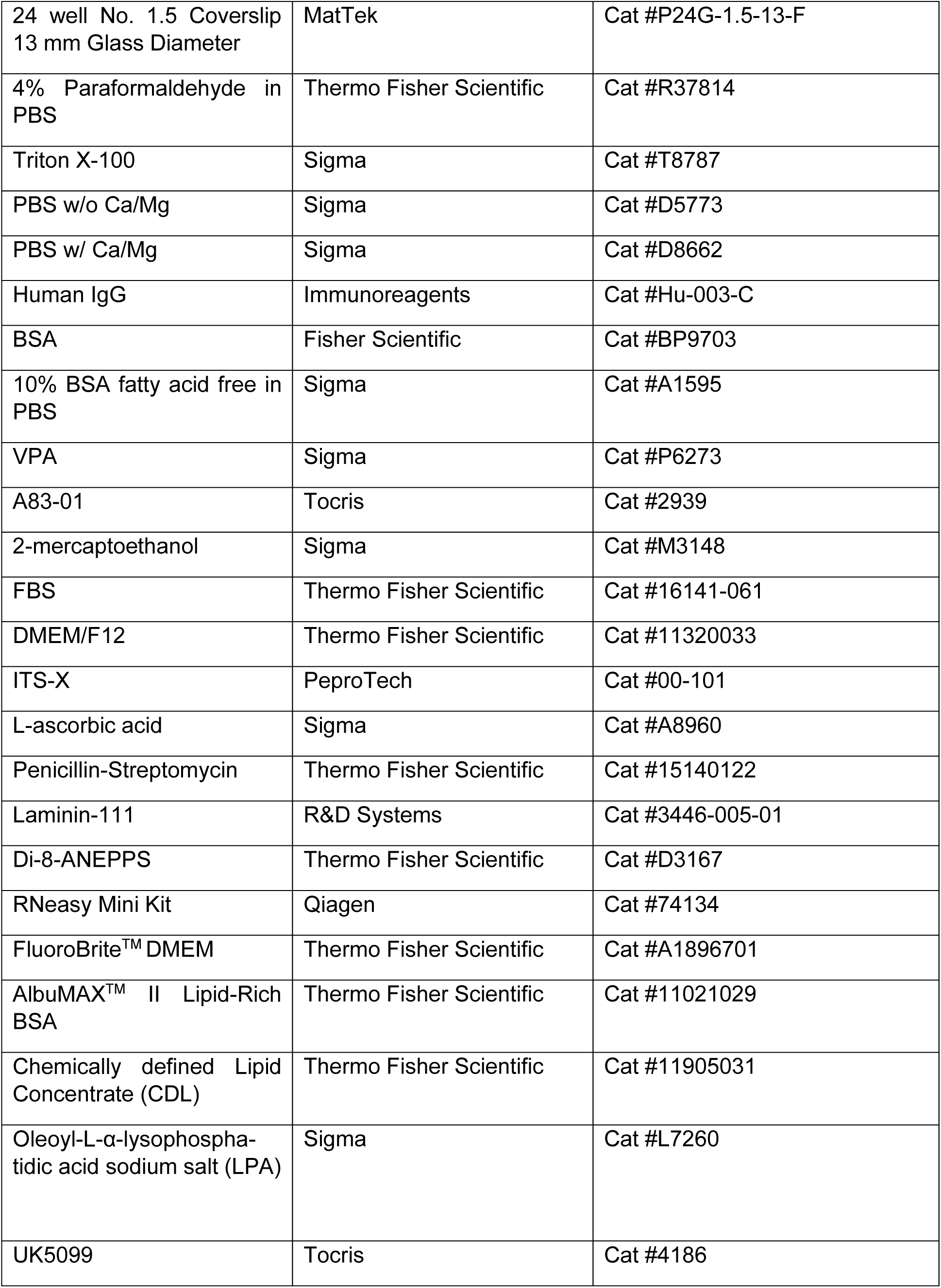
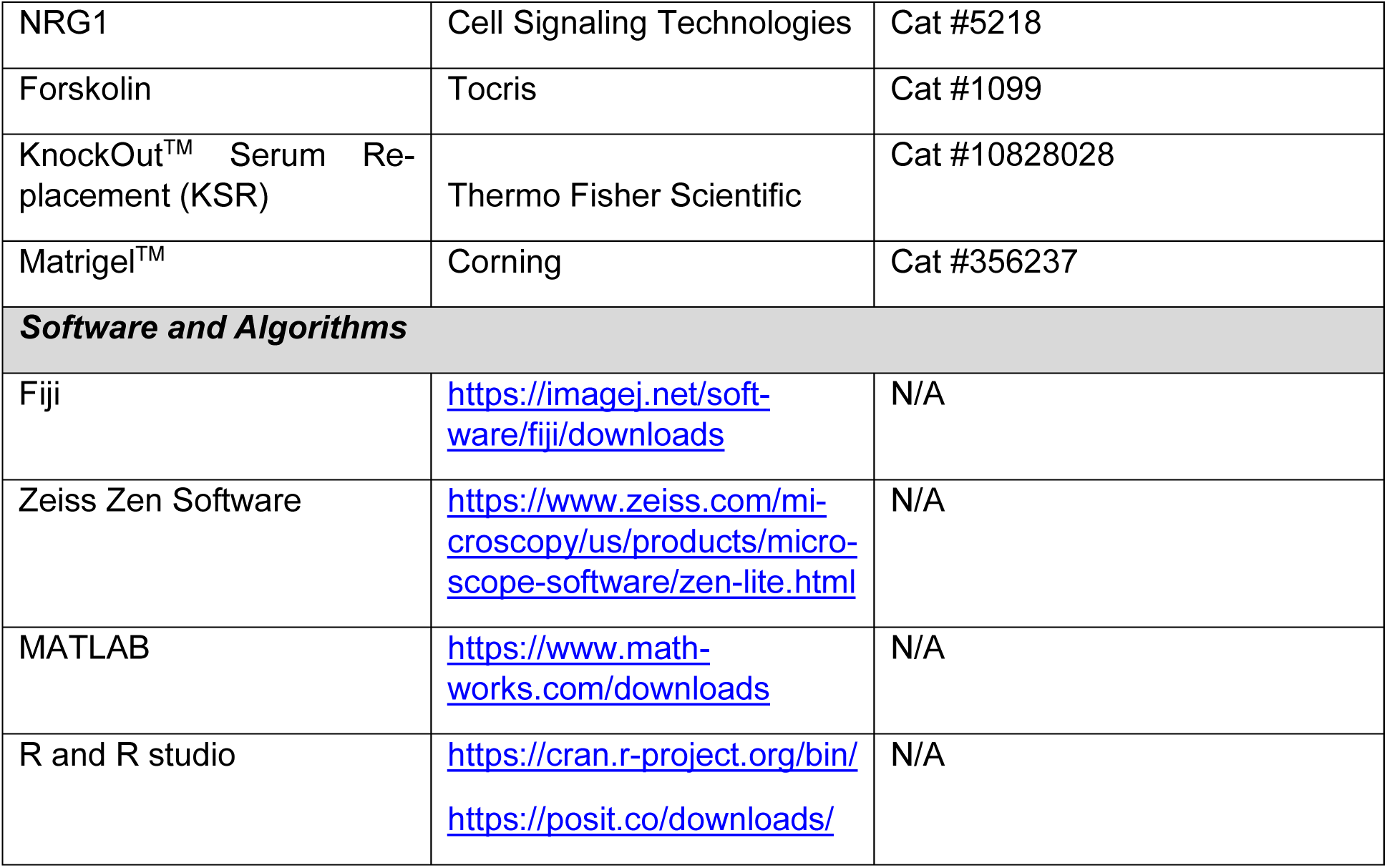
Key Resources.

### Derivation and culture of hTSCs

CTBs from two de-identified term placentas with no known pathology were provided by Amnion Foundation on a fee-for-service basis. Briefly, placental CTBs were enriched from placental tissue by Percoll gradient centrifugation and CD49f+ (i.e. integrin α6+) cells were subsequently isolated using standard protocols and frozen in aliquots of 500,000 cells each. Frozen aliquots of CTBs received from Amnion Foundation were thawed directly into wells of a 24-well plate pre-coated with 3 µg/ml of vitronectin and 1µg/mL of laminin-521, in TUA medium. TUA medium is trophoblast stem cell medium (TSCM) described by Okae et al. (5) (Dulbecco’s Modified Eagle Medium/Nutrient Mixture F-12 (DMEM/F-12) supplemented with 0.1 mM 2-mercaptoethanol, 0.2% FBS, 0.5% Penicillin-Streptomycin, 0.3% BSA, 1% Insulin-Transferrin-Selenium-Ethanolamine (ITS-X), 1.5 µg/mL L-ascorbic acid, 50 ng/mL EGF, 2 μM CHIR99021, 0.5 μM A83-01, 1 μM SB431542, 0.8 mM VPA, and 5 μM Y-27632), supplemented with 50 nM UK5099 and 0.2% AlbuMAX II. Cells were cultured in 5% CO_2_ at 37°C and medium was changed every other day. Initially, very slow cell growth was observed. Cells were passaged using treatment with TrypLE Express for 10-15 minutes at 37°C once cells became confluent or after 14 days, whichever was sooner; initial split ratios were 1:1-1:2. Subsequently, cells were transferred to 35 mm plates and routinely passaged every 7-10 days at a typical split ratio of 1:10. In some experiments, primary CTBs were also thawed directly into TSCM, TA medium (TSCM+0.2% AlbuMAX II), TUL medium (TSCM+50nM UK-5099+10 nM LPA), or TUC medium (TSCM+50nM UK-5099+0.1% chemically defined lipid concentrate). After 5 passages, T1 and T2 hTSCs derived in TUA medium were additionally transitioned into TSCM. T1 and T2 hTSCs, as well as CT29 and CT30 hTSCs derived from first trimester CTBs were cultured in TSCM as previously described (15).

### EVT and STB differentiation

EVT and STB differentiation was carried out using protocols previously described by Karakis et al.(15) or Okae et al (5). Prior to differentiation, hTSCs at confluence were dissociated into single cells using TrypLE Express, and cells were seeded into wells of a 24 well glass bottom plate (∼ 45,000) or 35 mm glass bottom plates; plates were pre-coated with 3 µg/ml of vitronectin and 1 µg/ml of Laminin-521. Studies using the protocol by Karakis et al. were conducted as follows: For STB differentiation, cells were seeded in defined trophoblast differentiation medium (DTDM; DMEM/F-12 with 1% ITS-X, 75 µg/mL L-ascorbic acid) supplemented with 50ng/mL EGF and 5 µM Y-27632 at passage. On day 2 and day 4, medium was replaced with DTDM. Cells were fixed on day 6. For EVT differentiation, cells were seeded in DTDM supplemented with 50 ng/mL EGF, 5 µM Y-27632, and 150 µg/mL laminin-111. On day 2 and day 4, medium was replaced with DTDM; cultures were analyzed on day 6. Studies using the protocol by Okae et al. were conducted as follows: For STB differentiation, cells were seeded in STB medium (DMEM/F12 supplemented with 0.1 mM 2-mercaptoethanol, 0.5% Penicillin-Streptomycin, 0.3% BSA, 1% ITS-X supplement, 2.5 μM Y-27632, 2 μM forskolin, and 4% KSR). Medium was replaced on day 3 and cells were fixed on day 6. For EVT differentiation, cells were seeded in cultured in EVT medium (DMEM/F12 supplemented with 0.1 mM 2-mercaptoethanol, 0.5% Penicillin-Streptomycin, 0.3% BSA, 1% ITS-X supplement, 100 ng/ml NRG1, 7.5 μM A83-01, 2.5 μM Y27632, and 4% KSR); Matrigel was added to a final media concentration of 2% after suspending the cells in EVT medium. On day 3, the medium was replaced with the EVT medium without NRG1 and Matrigel was added to a final concentration of 0.5%. Cells were fixed on day 6.

### Immunostaining

Cells were fixed with 4% paraformaldehyde for 10 minutes, permeabilized with 0.5% Triton X-100 in PBS for 10 minutes, and subsequently incubated in blocking buffer (0.5% BSA, and 200 μM human IgG in PBS) for at least one hour. Cells were then incubated overnight at 4°C on a rocker with primary antibodies diluted in blocking buffer. Primary antibodies used were anti-HLA-G (1:250), anti-Notch1 (1:250), anti-hCG (1:50), anti-SDC-1 (1:500), anti-KRT7 (1:50), anti-p63 (1:50), anti-TEAD4 (1:50), anti-CDX2 (1:250), anti-GATA3 (1:500), anti-YAP (1:200), anti-TFAP2C (1:400). Corresponding isotype controls (rabbit monoclonal IgG, rabbit XP IgG, mouse IgG1, mouse IgG2a, and mouse IgG2b) were used at the same concentrations as the primary antibodies. Subsequently, cells were washed twice with PBS, and labeled with secondary antibodies (Alexa Fluor 488-Rabbit or Alexa Fluor 647-Mouse conjugated antibodies) and DAPI in blocking buffer for 0.5-1 hr. Finally, cells were washed twice in PBS and stored in PBS prior to imaging with a laser scanning confocal microscope (LSM880, Carl Zeiss, Germany).

### Confocal microscopy image analysis

Image analysis was conducted using an image processing algorithm created in MATLAB R2021a, as previously described (15, 31). Briefly, TIFF images were generated after background correction in Zeiss Zen Lite software, using settings for corresponding isotype images for all samples. The algorithm identifies cells using DAPI staining (blue channel) and determines relative average protein expression intensity corresponding to antibody staining by mapping the red/green channels to individual cells based on the blue channel. For analysis of nuclear expression, only green/red pixels overlapping with the blue/pixels were considered. Expression intensity was set to zero if the average intensity is lower than the average intensity of all cells in the isotype control image. Data was normalized by the average isotype control expression intensity. Analysis for each experimental condition used 14 images (7 from each of two biological replicates); 7 images were used for isotype controls. This was performed for 14 isotype control images and 14 experimental images (7 images for each of two replicates). Statistical analysis was conducted using the non-parametric Mann-Whitney U test, in Microsoft Excel as described (15). Results of this test are given as a p-value to compare differences in medians. Statistical significance was inferred at p<0.05.

### Membrane staining and calculation of fusion index for STB differentiation

Cells were washed with PBS (with Mg^2+^ and Ca^2+^) and incubated with 1-2 µM Di-8-ANEPPS and DAPI in PBS (with Mg^2+^ and Ca^2+^) on ice for ∼0.5-1 hour. Subsequently, cells were washed once with FluoroBrite^TM^ DMEM and imaged in FluoroBrite^TM^ DMEM or PBS (with Mg^2+^ and Ca^2+^) using a Keyence BZ-X810 system. The fusion index for each condition was calculated using two biological replicates per cell line, except for T2 hTSCs in TUA medium differentiated using the protocol by Karakis et al., where a single replicate was used. A total of 7 images were captured for each replicate. Single cells and fused cells were individually quantified from each image, and the number of nuclei in each fused cell was determined. The fusion index for a single replicate was calculated as (N-S)/T, where N is the number of nuclei in syncytia, S is the number of syncytia, and T is total count of nuclei in both fused and unfused cells from 7 images. Fused cells with n>=3 nuclei were considered as syncytia.

### Total RNA extraction and sequencing

RNA was isolated using Purelink RNA Mini Kit using the manufacturer’s protocol and quantified using a Nanodrop 1000 spectrophotometer (Thermo Scientific, Waltham, MA). RNA samples were collected from at least three independent replicates, at different passage numbers. Library preparation with polyA selection and Illumina sequencing was conducted at Azenta Life Sciences (South Plainfield, NJ, USA), as follows. RNA samples were quantified using Qubit 2.0 Fluorometer (Life Technologies, Carlsbad, CA, USA) and RNA integrity was checked using Agilent TapeStation 4200 (Agilent Technologies, Palo Alto, CA, USA). RNA sequencing libraries were prepared using the NEBNext Ultra II RNA Library Prep Kit for Illumina using manufacturer’s instructions (NEB, Ipswich, MA, USA). Briefly, mRNAs were initially enriched with oligo-dT beads. Enriched mRNAs were fragmented for 15 minutes at 94 °C. First strand and second strand cDNA were subsequently synthesized. cDNA fragments were end repaired and adenylated at 3’ ends, and universal adapters were ligated to cDNA fragments, followed by index addition and library enrichment by PCR with limited cycles. The sequencing library was validated on the Agilent TapeStation (Agilent Technologies, Palo Alto, CA, USA), and quantified by using Qubit 2.0 Fluorometer as well as by quantitative PCR (KAPA Biosystems, Wilmington, MA, USA). The sequencing libraries were multiplexed and clustered onto a flowcell on the Illumina HiSeq 4000 instrument according to manufacturer’s instructions. The samples were sequenced using a 2x150bp Paired End (PE) configuration. Image analysis and base calling were conducted by the NovaSeq Control Software (NCS). Raw sequence data (.bcl files) generated from Illumina HiSeq was converted into FASTQ files and de-multiplexed using Illumina bcl2fastq 2.20 software. One mismatch was allowed for index sequence identification.

### Analysis of RNA-Seq data

Raw FASTQ formatted sequence reads were imported into CLC Genomics Workbench (version 23.0.2 and 25.0, QIAGEN Digital Insights). The imported reads were trimmed based on quality score and ambiguous nucleotides were also removed using default software settings. The trimmed sequences were mapped to the reference genome, GRCh38.110 using the RNA-Seq analysis feature in CLC Genomics Workbench; default mapping and expression settings were used. For principal component analysis (PCA) and differentially expressed gene analysis (DEG), “PCA for RNA-Seq” and “Differential Expression for RNA-Seq” toolsets were used, respectively. For heat maps, Euclidean distance with complete linkage option was used; a fixed number of 10,000 features, with 5 or more counts in at least one sample were used as filtering options. The heatmaps were clustered based on metadata groups. For targeted gene analysis, |log_2_(fold change)| ≥1 in expression between comparison groups with a threshold false discovery rate (FDR) adjusted p-value < 0.05 was used. The following datasets from the literature were used for comparative analysis: CT29 and CT30 hTSCs in TSCM(8) (GSE137295); hTSCs derived from term CTB in TSCM at 1% oxygen concentration(14) (GSE158901); primary vCTBs from the first trimester (6) (GSE109976). Additionally, the following datasets were used for comparative analysis with hTSCs from hPSCs: from Dong et al. (9) (GSE138762), we included GSM4116158 (H9 naïve TSC Bulk RNA-seq) and GSM4116160 (WIBR3 naïve TSC Bulk RNA-seq). From Guo et al. (22) (GSE167089), we included GSM5092604, GSM5092605 (hNES1), GSM5092608, GSM5092609 (niPSC2), and GSM5092612, GSM5092613 (niPSC4). From Io et al. (23) (GSE144994), we included GSM4304002, GSM4304003, GSM4304004, GSM4304005, GSM4304006, and GSM4304007. From Zorzan et al. (24) (GSE184562), we included GSM6754200, GSM6754201, and GSM6754202, representing replicates of ccTSC.02. From Mischler et al. (8) (GSE137295), we included GSM4074861, GSM4074862, and GSM4074863 (hTSCs from H9 hESCs in TSCM), and GSM4074865, GSM4074866, and GSM4074867 (hTSCs from H1 hESCs in TSCM). Unless otherwise specified, first-trimester hTSCs, CT 29 and CT 30 were used as the control group for analysis.

The list of differentially expressed genes for various conditions was imported into Ingenuity Pathway Analysis (IPA, Fall 2023 release version, QIAGEN Digital Insights) to analyze upregulated and downregulated biological pathways. A more stringent parameter list was used for IPA analysis to identify the most abundant pathways across conditions; filtering parameters were as follows: FDR p-value ≤0.05, Max group mean ≥ 5, |log_2_ fold-change| ≥ 2. Statistical |Z-score| cut-off was set at 1 with a - log_10_ p-value cutoff = 1.3 (p-value = 0.05).

### Cluster analysis and gene ontology (GO) for analysis of RNA-seq data

K-means cluster analysis was performed using R version 4.3.1 and R studio. Data of DEGs for different derivation conditions for hTSCs from term CTBs were collected from CLC genomic workbench and imported into R. The TPM values for these genes from multiple replicates were averaged and used for pairwise analysis. CT29 and CT30 TPM (transcripts per million) values were used as control in this pairwise analysis for these three term-derived hTSC conditions. Data processing involved removing missing values, assigning gene names as identifiers, and log2 transformation of TPM values to normalize the data. The optimal number of clusters (k) was determined using the Elbow method, which plots the Total Within-Cluster Sum of Squares (WSS) vs k-values (from 2 to 10 in this analysis). The sharpest decline in the plot determines optimal k-value. We found k-value equal to 3 for TUA and TSCM f TUA. But for 1% oxygen condition, k-value was equal to 4.

Following clustering, gene ontology (GO) analysis was performed separately for each gene cluster to identify enriched biological processes (BP). Gene lists from each cluster were extracted and subjected to GO enrichment analysis. We used clusterprofiler package in R. The analysis focused on the biological processes (BP) category. Default p-value and q-value were used for enrichgo function. The full list of GO enrichment results are available in **Tables S7-19**. The code used for this cluster and GO analysis is available at https://github.com/mjabn/K-means-clustering

## Supporting information

Supporting information includes 9 supplementary figures and 29 supplementary tables.

## Data availability

RNA-Seq data generated in this study have been deposited in Gene Expression Omnibus (GEO; accession number GSE267112). Additionally, RNA-Seq data from the literature used in this study are available in GEO (accession numbers GSE137295, GSE158901, and GSE109976, GSE138762, GSE167089, GSE144994, GSE184562, GSE137295). The code used for image analysis can be found at https://doi.org/10.5281/zenodo.7700524. All other data is available within the article and its supplementary materials.

## Author contributions

VK and BMR conceptualization; VK, JWB, MJ investigation, formal analysis, data curation and writing original draft; JWB, MJ, BMR data visualization; BMR editing and writing revised draft; VK, ASM software; ASM resources; BMR, VK funding acquisition; BMR project administration. VK, JWB, and MJ contributed equally and as such reserve the right to list their name first while listing this publication on professional documents.

## References

1. James, J. L. (2016) Stem Cells and Pregnancy Disorders: From Pathological Mechanisms to Therapeutic Horizons. Semin Reprod Med. 34, 17–26

2. Goldman-Wohl, D., and Yagel, S. (2002) Regulation of trophoblast invasion: From normal implantation to pre-eclampsia. in Molecular and Cellular Endocrinology, pp. 233–238, 187, 233–238

3. Norwitz, E. R. (2006) Defective implantation and placentation: Laying the blueprint for pregnancy complications, Reproductive Healthcare Ltd, 10.1016/S1472-6483(10)60649-9

4. Chavatte-Palmer, P., and Tarrade, A. (2016) Placentation in different mammalian species. Annales d’Endocrinologie. 77, 67–74

5. Okae, H., Toh, H., Sato, T., Hiura, H., Takahashi, S., Shirane, K., Kabayama, Y., Suyama, M., Sasaki, H., and Arima, T. (2018) Derivation of Human Trophoblast Stem Cells. Cell stem cell. 22, 50–63.e6

6. Haider, S., Meinhardt, G., Saleh, L., Kunihs, V., Gamperl, M., Kaindl, U., Ellinger, A., Burkard, T. R., Fiala, C., Pollheimer, J., Mendjan, S., Latos, P. A., and Knöfler, M. (2018) Self-Renewing Trophoblast Organoids Recapitulate the Developmental Program of the Early Human Placenta. Stem Cell Reports. 11, 537–551

7. Turco, M. Y., Gardner, L., Kay, R. G., Hamilton, R. S., Prater, M., Hollinshead, M. S., McWhinnie, A., Esposito, L., Fernando, R., Skelton, H., Reimann, F., Gribble, F. M., Sharkey, A., Marsh, S. G. E., O’rahilly, S., Hemberger, M., Burton, G. J., and Moffett, A. (2018) Trophoblast organoids as a model for maternal–fetal interactions during human placentation, Nature Publishing Group, 10.1038/s41586-018-0753-3

8. Mischler, A., Karakis, V., Mahinthakumar, J., Carberry, C. K., Miguel, A. S., Rager, J. E., Fry, R. C., and Rao, B. M. (2021) Two distinct trophectoderm lineage stem cells from human pluripotent stem cells. Journal of Biological Chemistry. 296, 100386

9. Dong, C., Beltcheva, M., Gontarz, P., Zhang, B., Popli, P., Fischer, L. A., Khan, S. A., Park, K.-M., Yoon, E.-J., Xing, X., Kommagani, R., Wang, T., Solnica-Krezel, L., and Theunissen, T. W. (2020) Derivation of trophoblast stem cells from naïve human pluripotent stem cells. eLife. 10.7554/elife.52504

10. Cinkornpumin, J. K., Kwon, S. Y., Guo, Y., Hossain, I., Sirois, J., Russett, C. S., Tseng, H. W., Okae, H., Arima, T., Duchaine, T. F., Liu, W., and Pastor, W. A. (2020) Naive Human Embryonic Stem Cells Can Give Rise to Cells with a Trophoblast-like Transcriptome and Methylome. Stem Cell Reports. 15, 198–213

11. Castel, G., Meistermann, D., Bretin, B., Firmin, J., Blin, J., Loubersac, S., Bruneau, A., Chevolleau, S., Kilens, S., Chariau, C., Gaignerie, A., Francheteau, Q., Kagawa, H., Charpentier, E., Flippe, L., François--Campion, V., Haider, S., Dietrich, B., Knöfler, M., Arima, T., Bourdon, J., Rivron, N., Masson, D., Fournier, T., Okae, H., Fréour, T., and David, L. (2020) Induction of Human Trophoblast Stem Cells from Somatic Cells and Pluripotent Stem Cells. Cell Reports. 33, 108419

12. Liu, X., Ouyang, J. F., Rossello, F. J., Tan, J. P., Davidson, K. C., Valdes, D. S., Schröder, J., Sun, Y. B. Y., Chen, J., Knaupp, A. S., Sun, G., Chy, H. S., Huang, Z., Pflueger, J., Firas, J., Tano, V., Buckberry, S., Paynter, J. M., Larcombe, M. R., Poppe, D., Choo, X. Y., O’Brien, C. M., Pastor, W. A., Chen, D., Leichter, A. L., Naeem, H., Tripathi, P., Das, P. P., Grubman, A., Powell, D. R., Laslett, A. L., David, L., Nilsson, S. K., Clark, A. T., Lister, R., Nefzger, C. M., Martelotto, L. G., Rackham, O. J. L., and Polo, J. M. (2020) Reprogramming roadmap reveals route to human induced trophoblast stem cells. Nature. 586, 101–107

13. Yang, L., Semmes, E. C., Ovies, C., Megli, C., Permar, S., Gilner, J. B., and Coyne, C. B. (2022) Innate immune signaling in trophoblast and decidua organoids defines differential antiviral defenses at the maternal-fetal interface. eLife. 11, e79794

14. Wang, L.-J., Chen, C.-P., Lee, Y.-S., Ng, P.-S., Chang, G.-D., Pao, Y.-H., Lo, H.-F., Peng, C.-H., Cheong, M.-L., and Chen, H. (2022) Functional antagonism between ΔNp63α and GCM1 regulates human trophoblast stemness and differentiation. Nat Commun. 13, 1626

15. Karakis, V., Jabeen, M., Britt, J. W., Cordiner, A., Mischler, A., Li, F., San Miguel, A., and Rao, B. M. (2023) Laminin switches terminal differentiation fate of human trophoblast stem cells under chemically defined culture conditions. Journal of Biological Chemistry. 10.1016/j.jbc.2023.104650

16. Arutyunyan, A., Roberts, K., Troulé, K., Wong, F. C. K., Sheridan, M. A., Kats, I., Garcia-Alonso, L., Velten, B., Hoo, R., Ruiz-Morales, E. R., Sancho-Serra, C., Shilts, J., Handfield, L.-F., Marconato, L., Tuck, E., Gardner, L., Mazzeo, C. I., Li, Q., Kelava, I., Wright, G. J., Prigmore, E., Teichmann, S. A., Bayraktar, O. A., Moffett, A., Stegle, O., Turco, M. Y., and Vento-Tormo, R. (2023) Spatial multiomics map of trophoblast development in early pregnancy. Nature. 10.1038/s41586-023-05869-0

17. Zhang, Y., Thery, F., Wu, N. C., Luhmann, E. K., Dussurget, O., Foecke, M., Bredow, C., Jiménez-Fernández, D., Leandro, K., Beling, A., Knobeloch, K.-P., Impens, F., Cossart, P., and Radoshevich, L. (2019) The in vivo ISGylome links ISG15 to metabolic pathways and autophagy upon Listeria monocytogenes infection. Nat Commun. 10, 5383

18. Albert, M., Vázquez, J., Falcón-Pérez, J. M., Balboa, M. A., Liesa, M., Balsinde, J., and Guerra, S. (2022) ISG15 Is a Novel Regulator of Lipid Metabolism during Vaccinia Virus Infection. Microbiology Spectrum. 10, e03893–22

19. Alcalá, S., Sancho, P., Martinelli, P., Navarro, D., Pedrero, C., Martín-Hijano, L., Valle, S., Earl, J., Rodríguez-Serrano, M., Ruiz-Cañas, L., Rojas, K., Carrato, A., García-Bermejo, L., Fernández-Moreno, M. Á., Hermann, P. C., and Bruno Sainz, J. (2020) ISG15 and ISGylation is required for pancreatic cancer stem cell mitophagy and metabolic plasticity. Nature Communications. 10.1038/s41467-020-16395-2

20. Bredow, C., Thery, F., Wirth, E. K., Ochs, S., Kespohl, M., Kleinau, G., Kelm, N., Gimber, N., Schmoranzer, J., Voss, M., Klingel, K., Spranger, J., Renko, K., Ralser, M., Mülleder, M., Heuser, A., Knobeloch, K.-P., Scheerer, P., Kirwan, J., Brüning, U., Berndt, N., Impens, F., and Beling, A. (2024) ISG15 blocks cardiac glycolysis and ensures sufficient mitochondrial energy production during Coxsackievirus B3 infection. Cardiovascular Research. 10.1093/cvr/cvae026

21. Bartho, L. A., O’Callaghan, J. L., Fisher, J. J., Cuffe, J. S. M., Kaitu’u-Lino, T. J., Hannan, N. J., Clifton, V. L., and Perkins, A. V. (2021) Analysis of mitochondrial regulatory transcripts in publicly available datasets with validation in placentae from pre-term, post-term and fetal growth restriction pregnancies. Placenta. 112, 162–171

22. Guo, G., Stirparo, G. G., Strawbridge, S. E., Spindlow, D., Yang, J., Clarke, J., Dattani, A., Yanagida, A., Li, M. A., Myers, S., Özel, B. N., Nichols, J., and Smith, A. (2021) Human naive epiblast cells possess unrestricted lineage potential. Cell Stem Cell. 28, 1040–1056.e6

23. Io, S., Kabata, M., Iemura, Y., Semi, K., Morone, N., Minagawa, A., Wang, B., Okamoto, I., Nakamura, T., Kojima, Y., Iwatani, C., Tsuchiya, H., Kaswandy, B., Kondoh, E., Kaneko, S., Woltjen, K., Saitou, M., Yamamoto, T., Mandai, M., and Takashima, Y. (2021) Capturing human trophoblast development with naive pluripotent stem cells in vitro. Cell Stem Cell. 28, 1023–1039.e13

24. Zorzan, I., Betto, R. M., Rossignoli, G., Arboit, M., Drusin, A., Corridori, C., Martini, P., and Martello, G. (2023) Chemical conversion of human conventional PSCs to TSCs following transient naive gene activation. EMBO reports. 24, e55235

25. Xu, F., Deng, C., Ren, Z., Sun, L., Meng, Y., Liu, W., Wan, J., and Chen, G. (2021) Lyso-phosphatidic acid shifts metabolic and transcriptional landscapes to induce a distinct cellular state in human pluripotent stem cells. Cell Reports. 37, 110063

26. Mylonis, I., Simos, G., and Paraskeva, E. (2019) Hypoxia-Inducible Factors and the Regulation of Lipid Metabolism. Cells. 8, 214

27. Mihaylova, M. M., Cheng, C.-W., Cao, A. Q., Tripathi, S., Mana, M. D., Bauer-Rowe, K. E., Abu-Remaileh, M., Clavain, L., Erdemir, A., Lewis, C. A., Freinkman, E., Dickey, A. S., La Spada, A. R., Huang, Y., Bell, G. W., Deshpande, V., Carmeliet, P., Katajisto, P., Sabatini, D. M., and Yilmaz, Ö. H. (2018) Fasting Activates Fatty Acid Oxidation to Enhance Intestinal Stem Cell Function during Homeostasis and Aging. Cell Stem Cell. 22, 769–778.e4

28. Veliova, M., Ferreira, C. M., Benador, I. Y., Jones, A. E., Mahdaviani, K., Brownstein, A. J., Desousa, B. R., Acín-Pérez, R., Petcherski, A., Assali, E. A., Stiles, L., Divakaruni, A. S., Prentki, M., Corkey, B. E., Liesa, M., Oliveira, M. F., and Shirihai, O. S. (2020) Blocking mitochondrial pyruvate import in brown adipocytes induces energy wasting via lipid cycling. EMBO Rep. 21, e49634

29. Ozmen, A., Guzeloglu-Kayisli, O., Tabak, S., Guo, X., Semerci, N., Nwabuobi, C., Larsen, K., Wells, A., Uyar, A., Arlier, S., Wickramage, I., Alhasan, H., Totary-Jain, H., Schatz, F., Odibo, A. O., Lockwood, C. J., and Kayisli, U. A. (2022) Preeclampsia is Associated With Reduced ISG15 Levels Impairing Extravillous Trophoblast Invasion. Front Cell Dev Biol. 10, 898088

30. Tecalco-Cruz, A. C., and Zepeda-Cervantes, J. (2023) Protein ISGylation: a posttranslational modification with implications for malignant neoplasms. Explor Target Antitumor Ther. 4, 699–715

31. Jabeen, M., Karakis, V., Britt, J., Miguel, A. S., and Rao, B. (2024) A quantitative image analysis platform for assessing trophoblast differentiation. Placenta. 10.1016/j.placenta.2024.07.009

