## Supporting Information for "Derivation of human trophoblast stem cells from placentas at birth"

[+]These authors contributed equally.

**Contents**

Figures S1-S9

Tables S1-S29 (in .xlsx format)

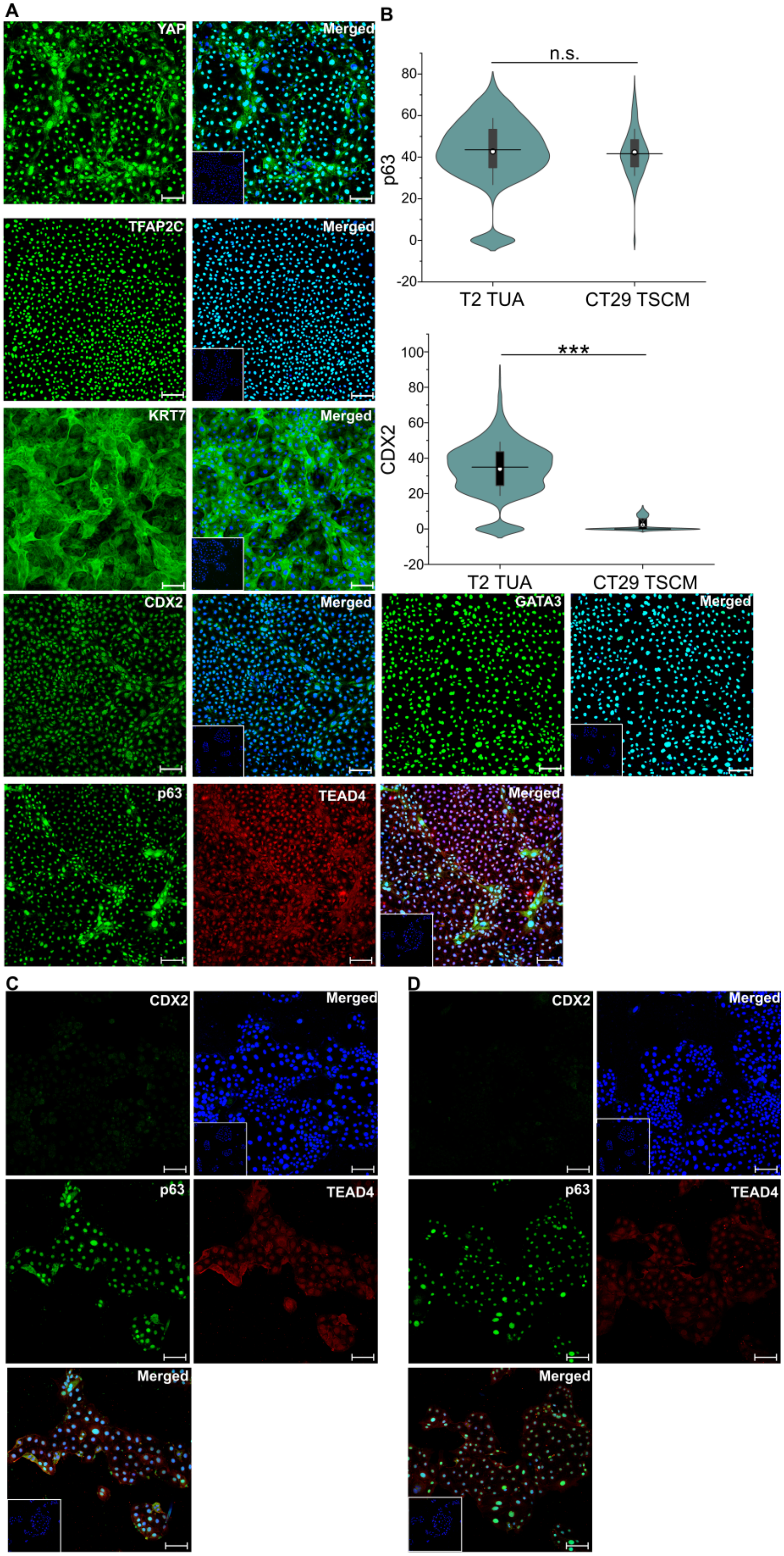
A
YAP
Merged
TFAP2C
Merged
KRT7
Merged
CDX2
Merged
GATA3
Merged
p63
TEAD4
Merged
B
n.s.
p63
T2 TUA
CT29 TSCM
***
CDX2
T2 TUA
CT29 TSCM
C
CDX2
Merged
p63
TEAD4
Merged
D
CDX2
Merged
p63
TEAD4
Merged

**Figure S1: Expression of hTSC markers in TUA medium.**

(A) Confocal microscopy imaging of T2 hTSCs cultured in TUA medium, staining for TFAP2C, YAP, GATA3, KRT7, TEAD4, p63, and CDX2; p63 was co-stained with TEAD4. Nuclei were stained with DAPI (blue). Isotype control is shown as an inset image.

(B) Quantification of expression of p63 and CDX2 from T2 hTSCs (n= 9025 for p63, n=9449 for CDX2) and CT29 hTSCs (n=2268 for p63, n=10259 for CDX2). Data from two biological replicates used. White circle represents the mean and the black line represents the median (***p-value < 0.001).

(C) Confocal microscopy imaging of CT29 hTSCs cultured in TSCM, staining for CDX2, p63 and TEAD4; p63 was co-stained with TEAD4. Nuclei were stained with DAPI (blue). Isotype control is shown as an inset image. Images for CDX2 and p63 staining are representative of those used for quantification of expression of CDX2 and p63 in **Fig. S1B**.

(D) Confocal microscopy imaging of CT30 hTSCs cultured in TSCM, staining for CDX2, p63 and TEAD4; p63 was co-stained with TEAD4. Nuclei were stained with DAPI (blue). Isotype control is shown as an inset image. Images for CDX2 and p63 staining are representative of those used for quantification of expression of CDX2 and p63 in **Fig. 1B**.

Scale bars are 100 μm for all images.

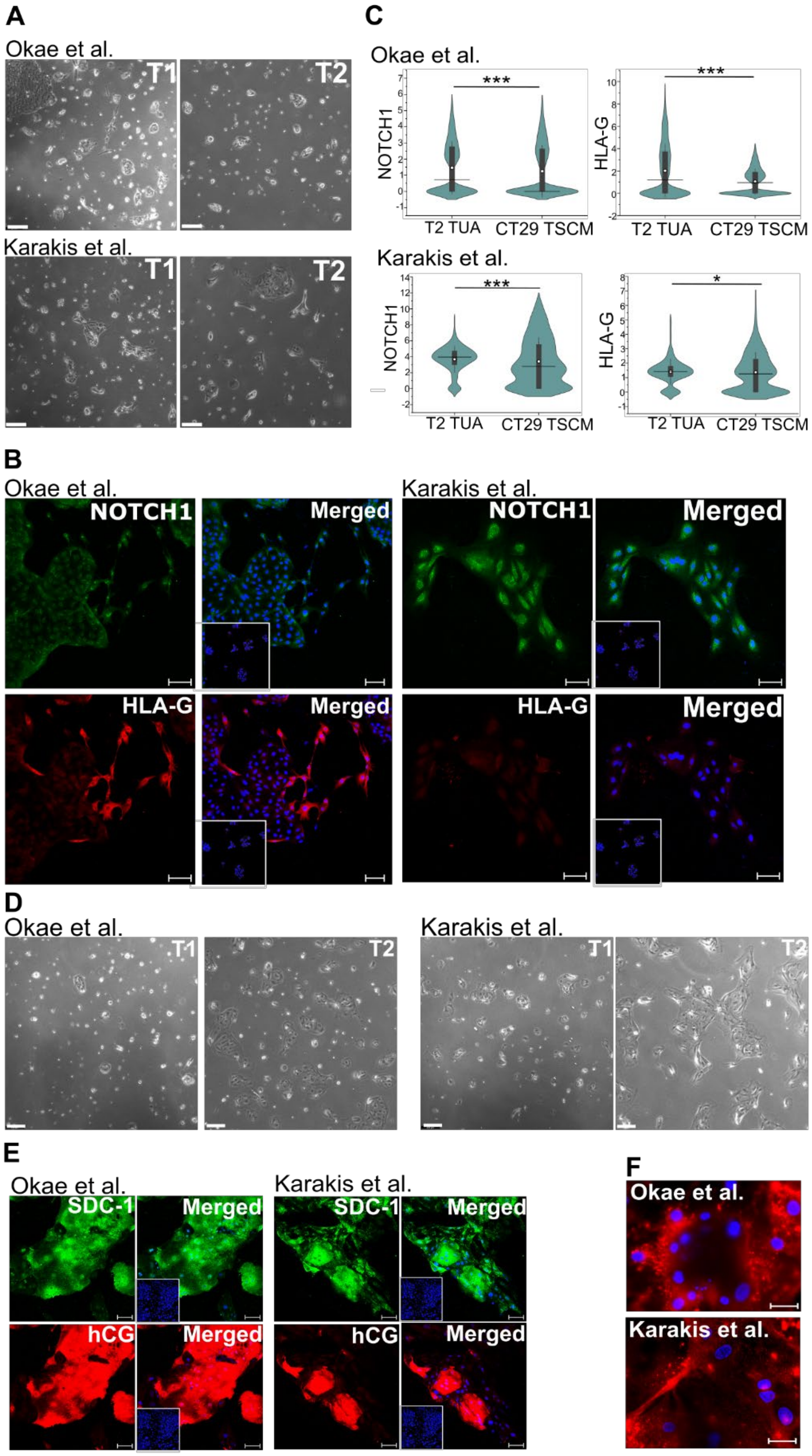
A
Okae et al.
T1
T2
Karakis et al.
T1
T2
C
Okae et al.
NOTCH1
***
T2 TUA
CT29 TSCM
HLA-G
***
T2 TUA
CT29 TSCM
Karakis et al.
NOTCH1
***
T2 TUA
CT29 TSCM
HLA-G
*
T2 TUA
CT29 TSCM
B
Okae et al.
NOTCH1
Merged
HLA-G
Merged
Karakis et al.
NOTCH1
Merged
HLA-G
Merged
D
Okae et al.
T1
T2
Karakis et al.
T1
T2
E
Okae et al.
SDC-1
Merged
hCG
Merged
Karakis et al.
SDC-1
Merged
hCG
Merged
F
Okae et al.
Karakis et al.

**Figure S2: EVT and STB differentiation of hTSCs from term CTBs in TUA medium.**

(A) Bright field images of EVT for T1 and T2 hTSCs differentiated to EVT using protocols by Okae et al. or Karakis et al.

(B) Confocal microscopy imaging of T2 hTSCs cultured in TUA and differentiated to EVTs using protocols by Okae et al. or Karakis et al., staining for NOTCH1 and HLA-G at day 6 of differentiation. Nuclei was stained with DAPI. Inset image is isotype control.

(C) Quantification of NOTCH1 and HLA-G expression from T2 cultured in TUA and CT29 hTSCs differentiated to EVTs using protocols by Okae et al. or Karakis et al. Data from two biological replicates used. White circle represents the mean and the black line represents the median. For T2 hTSCs using protocol by Okae et al., n=4922; Karakis et al., n=541. For CT29 hTSCs using protocol by Okae et al., n=6989; Karakis et al., n=807. (***p-value < 0.001, *p-value < 0.05)). Data for EVT differentiation using protocol by Okae et al. has been obtained from our previously published work (1) and re-analyzed for this figure.

(D) Bright field images of EVT for T1 and T2 hTSCs differentiated to STB using protocols by Okae et al. or Karakis et al.

(E) Confocal microscopy imaging staining for hCG and SDC-1 at day 6, for T2 hTSCs differentiated to STB using protocols by Okae et al. or Karakis et al. Nuclei were stained with DAPI. Inset images are isotype control.

(F) Di-8-ANEPPS membrane staining at day 6 for T2 hTSCs differentiated to STB using protocols by Okae et al. or Karakis et al.

Scale bars are 100 μm for confocal images and 250 μm for bright field images.

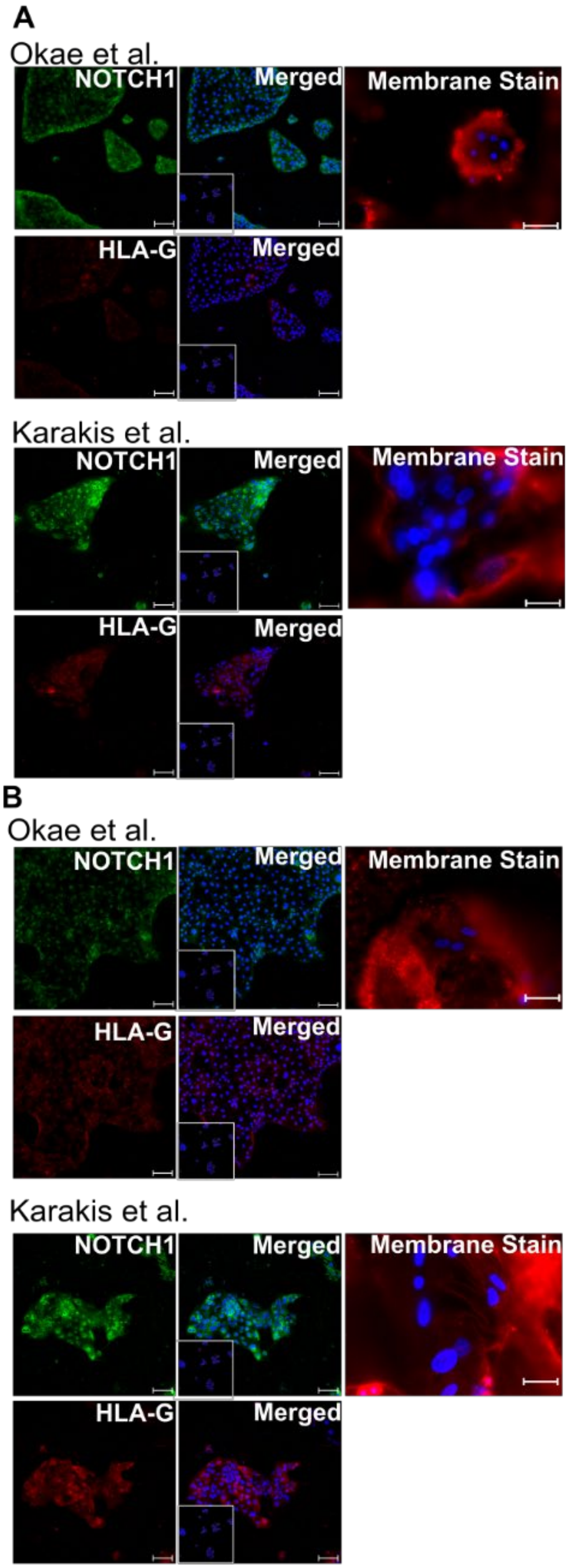
A
Okae et al.
NOTCH1
Merged
Membrane Stain
HLA-G
Merged
Karakis et al.
NOTCH1
Merged
Membrane Stain
HLA-G
Merged
B
Okae et al.
NOTCH1
Merged
Membrane Stain
HLA-G
Merged
Karakis et al.
NOTCH1
Merged
Membrane Stain
HLA-G
Merged

**Figure S3: EVT and STB differentiation of first trimester hTSCs.**

(A) Confocal microscopy imaging of CT29 hTSCs in TSCM differentiated to EVTs, staining for NOTCH1 and HLA-G at day 6, and Di-8-ANEPPS membrane staining for CT29 hTSCs differentiated to STB at day 6. Protocols by Okae et al. or Karakis et al. were used for hTSC differentiation. Images for NOTCH1 and HLA-G staining are representative of those used for quantification of NOTCH1 and HLA-G expression in **Fig. S2C**. Images for Di-8- ANEPPS membrane staining are representative of those used for calculation of fusion index in Fig. 2D. Data for EVT and STB differentiation using protocol by Okae et al. have been taken from our previously published work (1).

(B) Confocal microscopy imaging of CT30 hTSCs in TSCM differentiated to EVTs, staining for NOTCH1 and HLA-G, and Di-8-ANEPPS membrane staining for CT30 hTSCs differentiated to STB. Protocols by Okae et al. or Karakis et al. were used for hTSC differentiation. Images for NOTCH1 and HLA-G staining are representative of those used for quantification of NOTCH1 and HLA-G expression in **Fig. 2B**. Images for Di-8-ANEPPS membrane staining are representative of those used for calculation of fusion index in **Fig. 2D**. Data for EVT and STB differentiation using protocol by Okae et al. have been taken from our previously published work.

Scale bars are 100 μm for all images.

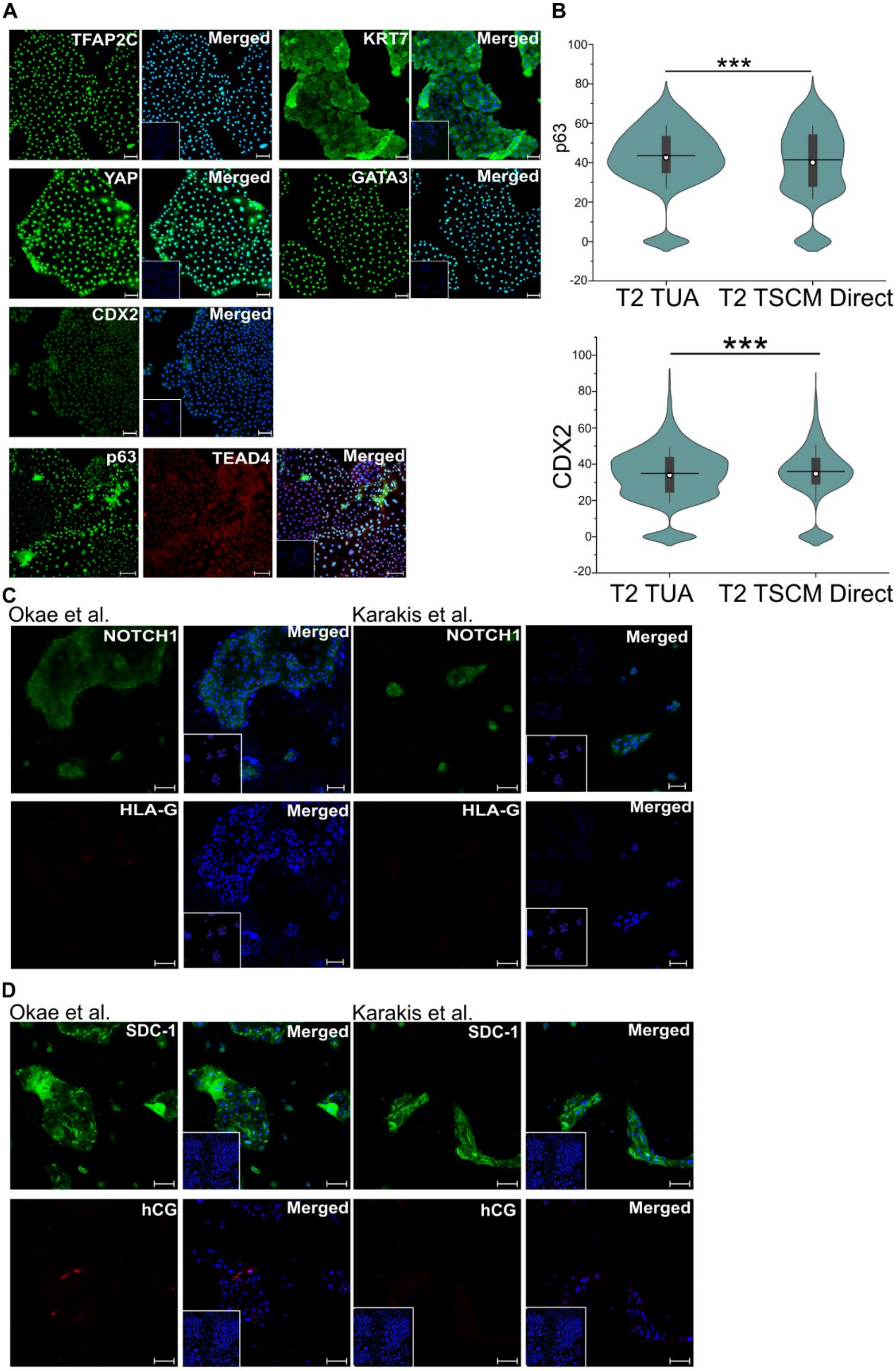
A
TFAP2C
Merged
KRT7
Merged
YAP
Merged
GATA3
Merged
CDX2
Merged
p63
TEAD4
Merged
B
***
p63
T2 TUA
T2 TSCM Direct
***
CDX2
T2 TUA
T2 TSCM Direct
C
Okae et al.
NOTCH1
Merged
Karakis et al.
NOTCH1
Merged
HLA-G
Merged
HLA-G
Merged
D
Okae et al.
SDC-1
Merged
Karakis et al.
SDC-1
Merged
hCG
Merged
hCG
Merged

**Figure S4: Attempted derivation of hTSCs from term CTBs in TSCM.**

(A) Confocal images of cells obtained during attempted derivation of hTSCs from term CTBs in TSCM, staining for TFAP2C, YAP, GATA3, KRT7, p63, TEAD4, and CDX2. Nuclei were stained with DAPI. Inset images are isotype controls. Primary CTBs are from the same placenta as those used for deriving T2 hTSCs in TUA medium.

(B) Quantification of expression of p63 and CDX2 in cells obtained during attempted derivation of hTSCs from term CTBs in TSCM (labeled TSCM direct). Data from two biological replicates; data from T2 hTSCs in TUA medium (same data as in **Fig. 1**) is shown for comparison. For TSCM direct, n=5043 for p63 staining and n=6046 for CDX2 staining. Data from two biological replicates used: data from T2 hTSCs in TUA medium (same data as in **Fig. S1**). White circle represents the mean and the black line represents the median (***p-value < 0.001).

(C) Confocal microscopy imaging of cells obtained during attempted derivation of hTSCs, differentiated to EVTs for 6 days using protocols by Okae et al. or Karakis et al., staining for NOTCH1 and HLA-G. Nuclei were stained with DAPI (blue). Inset images are isotype control.

(D) Confocal microscopy imaging of cells obtained during attempted derivation of hTSCs, differentiated to STB for 6 days using protocols by Okae et al. or Karakis et al., staining for SDC1 and hCG. Nuclei were stained with DAPI (blue). Inset images are isotype control.

Scale bars are 100 μm for all images.

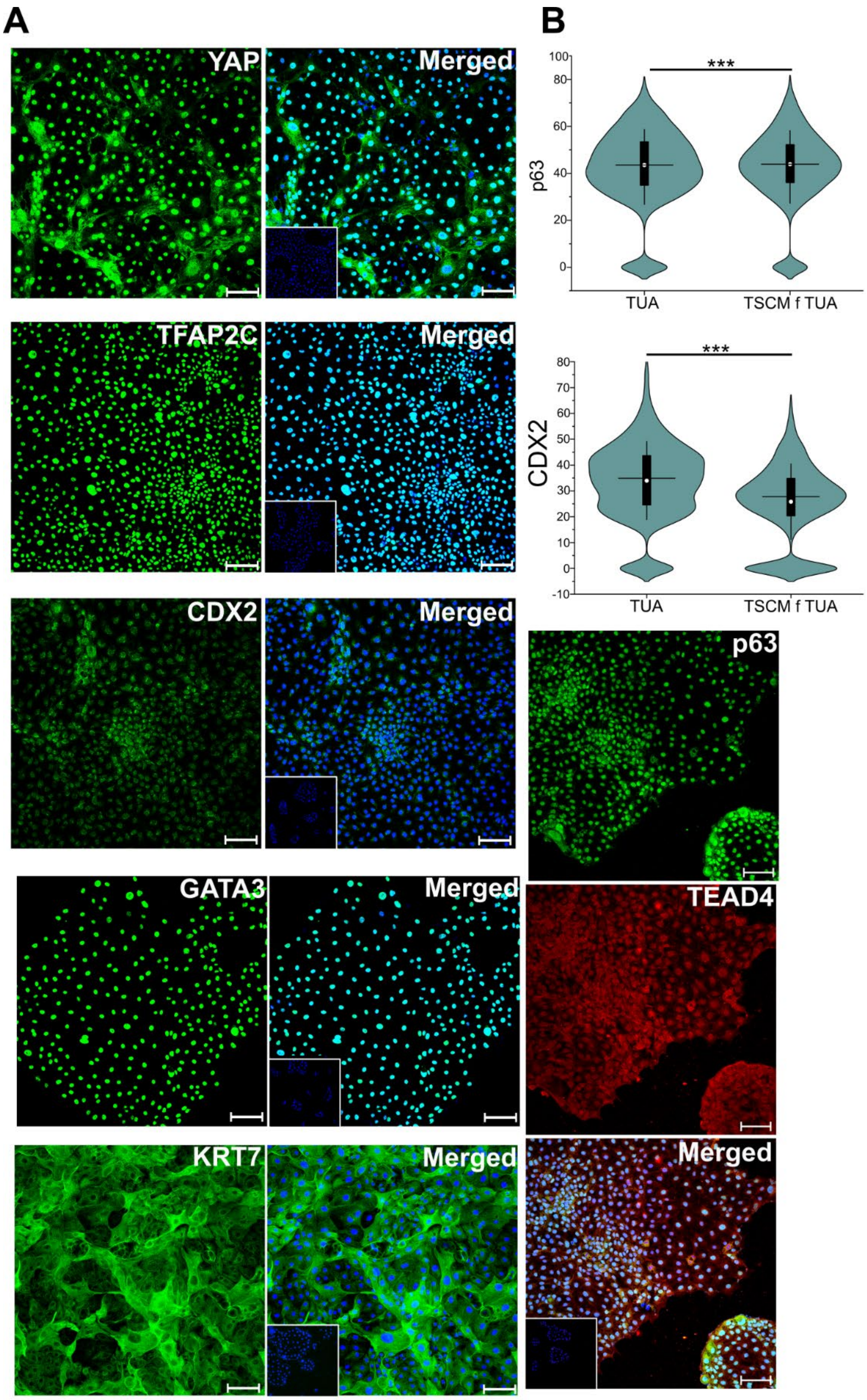
A
B
YAP
Merged
TFAP2C
Merged
CDX2
Merged
GATA3
Merged
KRT7
Merged
p63
TEAD4
Merged
***
p63
TUA
TSCM f TUA
***
CDX2
TUA
TSCM f TUA

**Figure S5: Expression of hTSC markers in cells transitioned to TSCM.**

(A) Confocal microscopy imaging of T2 hTSCs transitioned to TSCM, staining for TFAP2C, YAP, GATA3, KRT7, p63, TEAD4, and CDX2; p63 was co-stained with TEAD4. Nuclei were stained with DAPI. Inset images are isotype control. Scale bars are 100 μm.

(B) Quantification of expression of p63 and CDX2 expression from T2 hTSCs transitioned into TSCM (TSCM f TUA; for p63 n=7928, for CDX2 n= 9026). Data from two biological replicates used; data from T2 hTSCs in TUA medium (same data as in **Fig. S1**) is shown for comparison. White circle represents the mean and the black line represents the median (***p-value < 0.001).

Scale bars are 100 μm for all images unless specified otherwise.

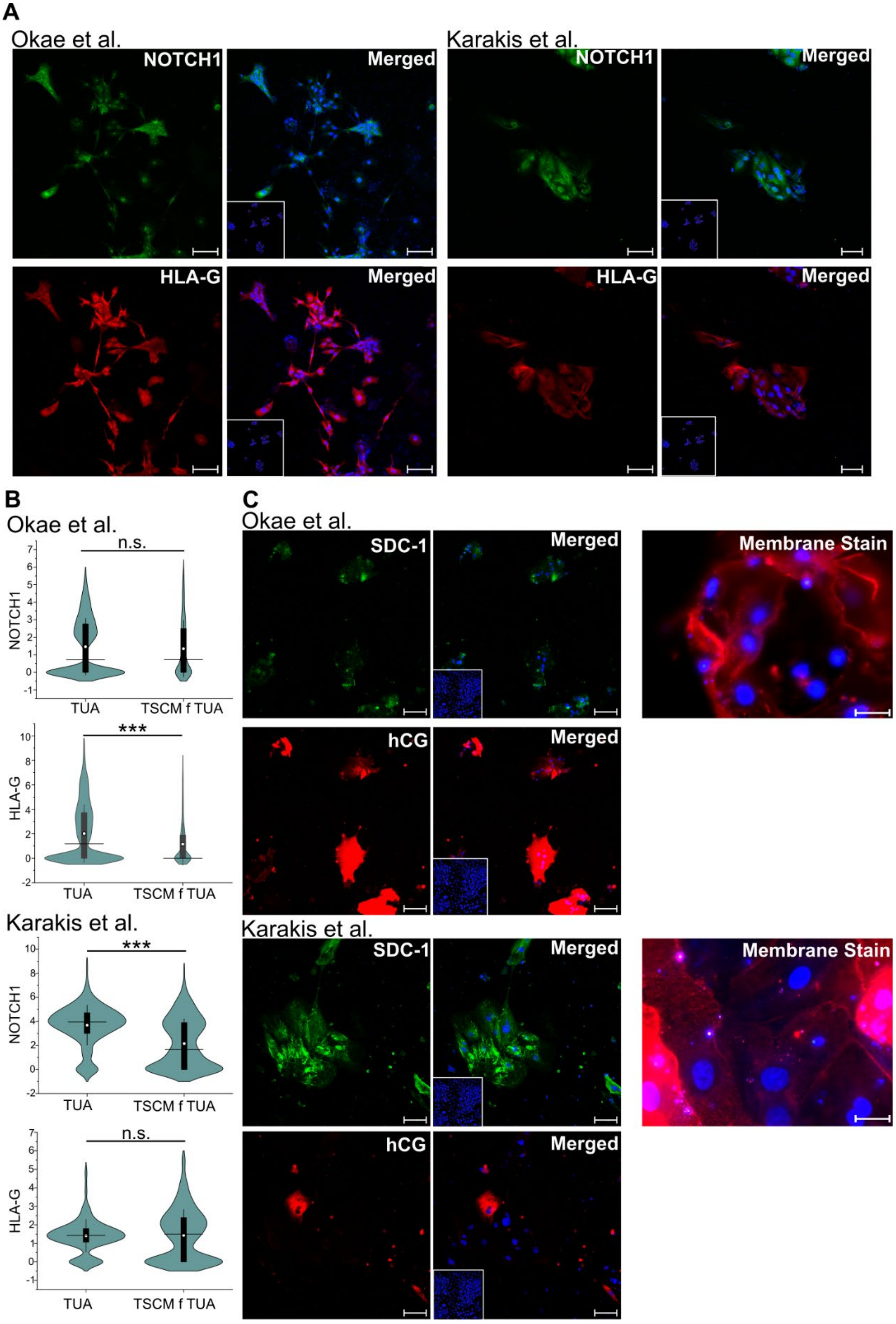
A
Okae et al.
NOTCH1
Merged
HLA-G
Merged
Karakis et al.
NOTCH1
Merged
HLA-G
Merged
B
Okae et al.
n.s.
NOTCH1
TUA
TSCM f TUA
***
HLA-G
TUA
TSCM f TUA
Karakis et al.
***
NOTCH1
TUA
TSCM f TUA
n.s.
HLA-G
TUA
TSCM f TUA
C
Okae et al.
SDC-1
Merged
Membrane Stain
hCG
Merged
Karakis et al.
SDC-1
Merged
Membrane Stain
hCG
Merged

**Figure S6: EVT and STB differentiation of hTSCs transitioned into TSCM.**

(A) Confocal microscopy imaging of T2 hTSCs transitioned into TSCM and differentiated for 6 days using protocols by Okae et al. or Karakis et al., staining for NOTCH1 and HLA-G at day 6 of differentiation. Nuclei were stained with DAPI.

(B) Quantification of NOTCH1 and HLA-G expression from T2 hTSCs transitioned into TSCM and differentiated with protocols by Okae et al. (n=1021) or Karakis et al. (n=540). Data from two biological replicates used. Data from T2 hTSCs in TUA medium is shown for comparison (same data as in **Fig. S2**). White circle represents the mean and the black line represents the median. (***p-value < 0.001, n.s. = not statistically significant (p>0.05)).

(C) Di-8-ANEPPS membrane staining and confocal microscopy imaging staining for hCG and SDC-1 at day 6, for T2 hTSCs in TSCM, differentiated to STB using protocols by Okae et al. or Karakis et al. Nuclei were stained with DAPI.

Scale bars are 100 μm for all images

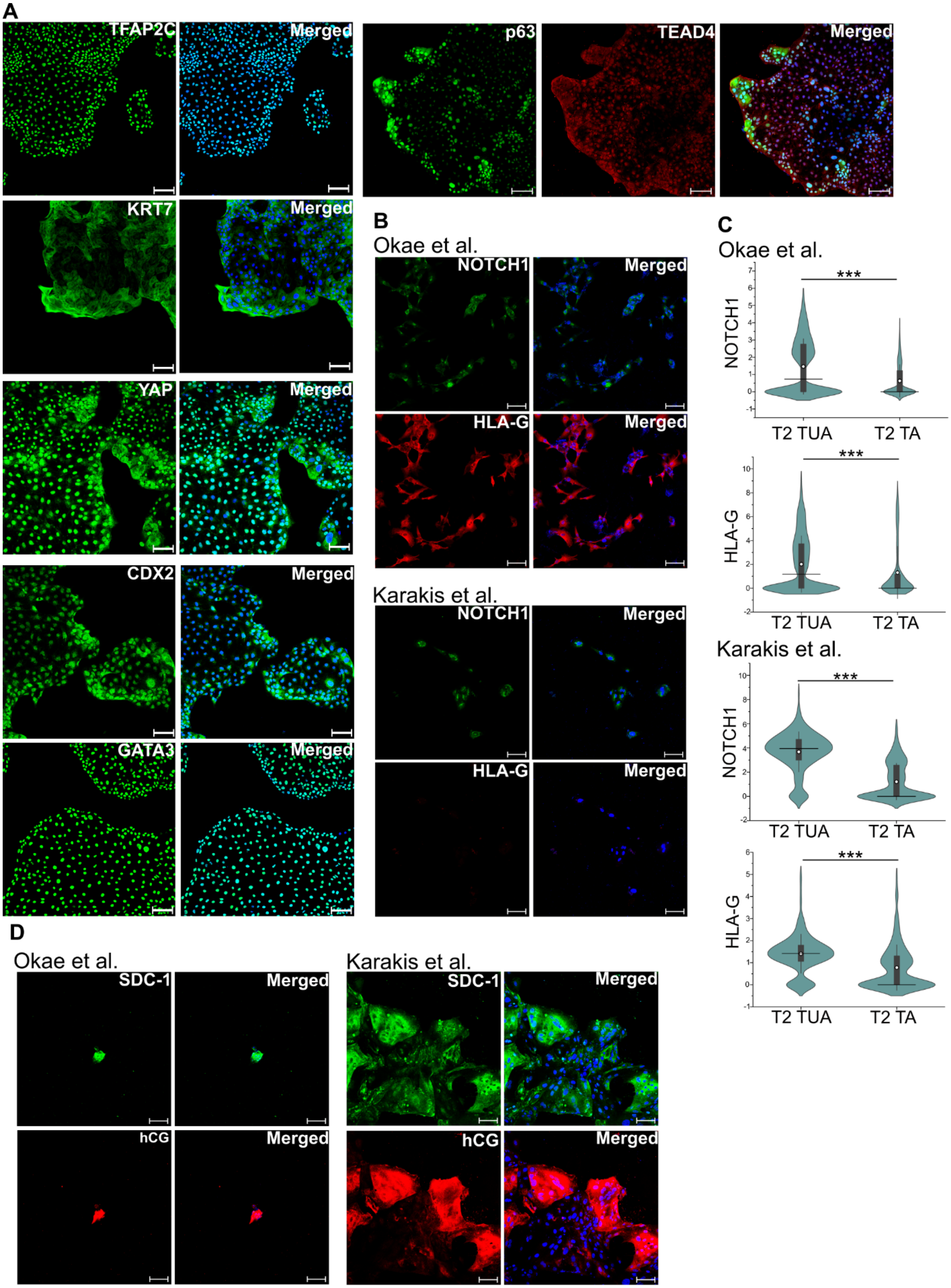
A
TFAP2C
Merged
p63
TEAD4
Merged
KRT7
Merged
YAP
Merged
CDX2
Merged
GATA3
Merged
B
Okae et al.
NOTCH1
Merged
HLA-G
Merged
Karakis et al.
NOTCH1
Merged
HLA-G
Merged
C
Okae et al.
NOTCH1
***
T2 TUA
T2 TA
HLA-G
***
T2 TUA
T2 TA
Karakis et al.
NOTCH1
***
T2 TUA
T2 TA
HLA-G
***
T2 TUA
T2 TA
D
Okae et al.
SDC-1
Merged
hCG
Merged
Karakis et al.
SDC-1
Merged
hCG
Merged

**Figure S7: Attempted derivation of hTSCs from term CTBs in TA medium.**

(A) Confocal imaging of cells obtained during attempted derivation of hTSCs in TA medium, staining for TFAP2C, YAP, GATA3, KRT7, p63, TEAD4, and CDX2; p63 was co-stained with TEAD4. Nuclei were stained with DAPI. Primary CTBs are from the same placenta as those used for deriving T2 hTSCs in TUA medium.

(B) Confocal microscopy imaging of cells obtained during attempted derivation of hTSCs in TA medium, differentiated to EVTs for 6 days using protocols by Okae et al. or Karakis et al., staining for NOTCH1 and HLA-G. Nuclei were stained with DAPI (blue).

(C) Quantification of NOTCH1 and HLA-G expression in cells obtained during attempted derivation of term CTBs in TA medium (labeled T2 TA), differentiated with protocols by Okae et al. (n=2058) or Karakis et al. (n=626). Data from two biological replicates used. Data from T2 hTSCs in TUA medium is shown for comparison (same data as in **Fig. S2**). White circle represents the mean and the black line represents the median. (*p-value < 0.05, **p-value < 0.001)

(D) Confocal microscopy imaging of cells obtained during attempted derivation of hTSCs in TA medium, differentiated to STB for 6 days using protocols by Okae et al. or Karakis et al., staining for SDC-1 and hCG. Nuclei were stained with DAPI (blue).

Scale bars are 100μm for all images.

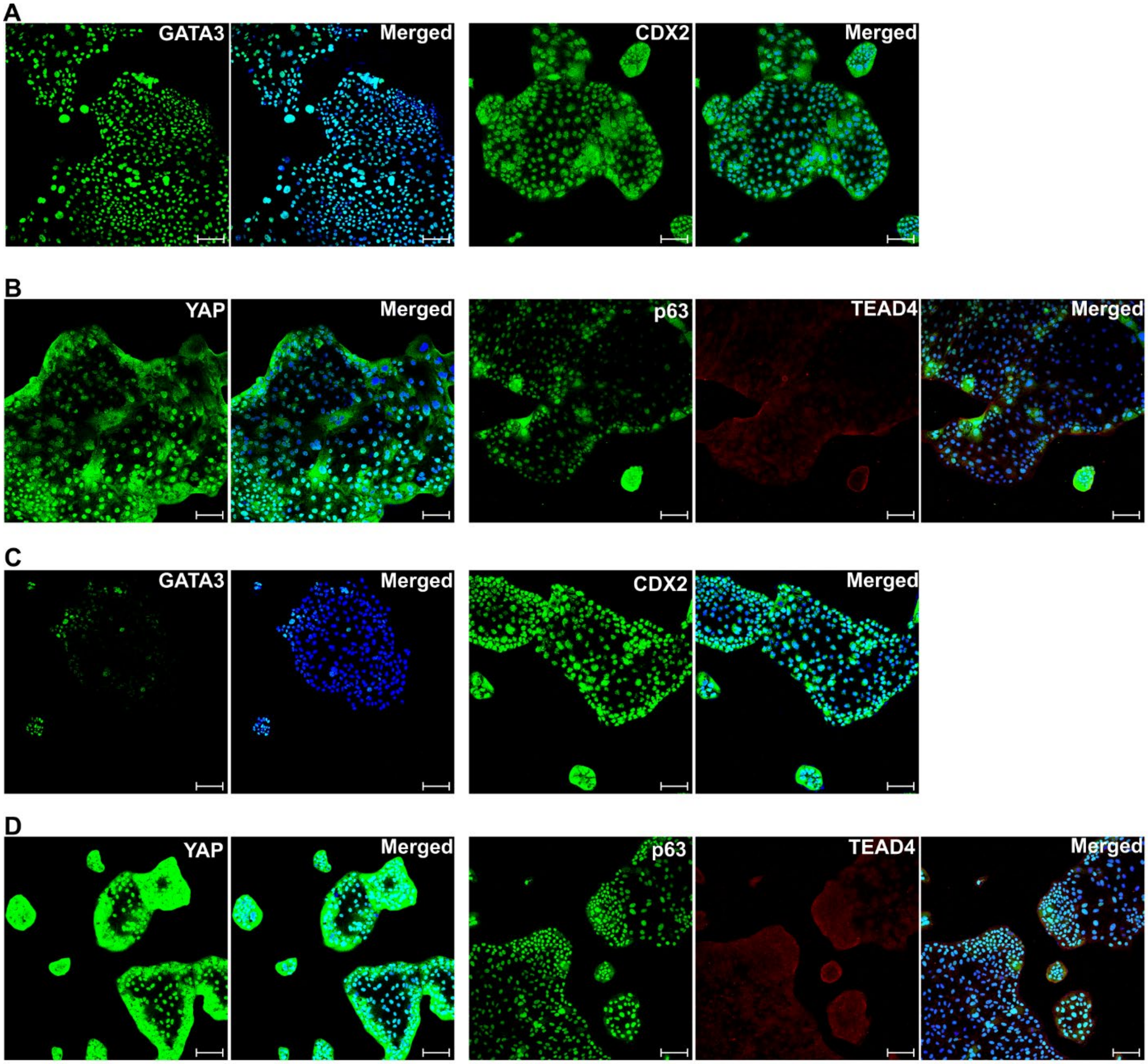


**Figure S8: Attempted derivation of hTSCs from term CTBs in TUL and TUC medium.**

Confocal microscopy imaging of cells obtained during attempted derivation of hTSCs in TUL medium (A-B) or TUC medium (C-D), staining for TFAP2C, YAP, GATA3, KRT7, p63, TEAD4, and CDX2; p63 was co-stained with TEAD4. Nuclei were stained with DAPI. Primary CTBs are from the same placenta as those used for deriving T1 hTSCs in TUA medium. Scale bars are 100μm for all images.

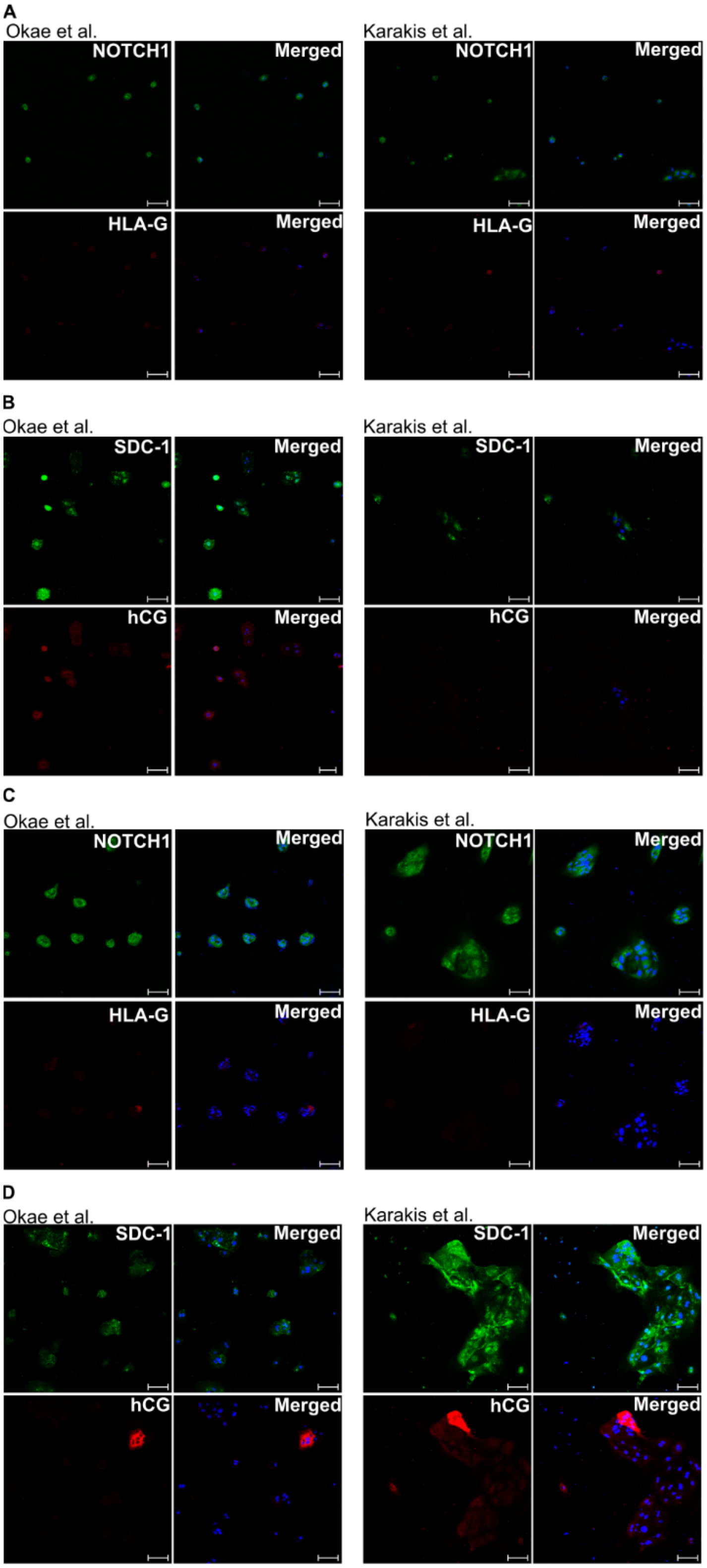
A
Okae et al.
NOTCH1
Merged
HLA-G
Merged
Karakis et al.
NOTCH1
Merged
HLA-G
Merged
B
Okae et al.
SDC-1
Merged
hCG
Merged
Karakis et al.
SDC-1
Merged
hCG
Merged
C
Okae et al.
NOTCH1
Merged
HLA-G
Merged
Karakis et al.
NOTCH1
Merged
HLA-G
Merged
D
Okae et al.
SDC-1
Merged
hCG
Merged
Karakis et al.
SDC-1
Merged
hCG
Merged

**Figure S9: Differentiation studies on cells obtained during attempted derivation of hTSCs derived in TUL and TUC medium.**

(A) Confocal microscopy imaging of cells obtained during attempted derivation of hTSCs in TUL medium, differentiated to EVTs for 6 days using protocols by Okae et al. or Karakis et al., staining for NOTCH1 and HLA-G. Primary CTBs are from the same placenta as those used for deriving T1 hTSCs in TUA medium.

(B) Confocal microscopy imaging of cells obtained during attempted derivation of hTSCs in TUL medium, differentiated to STB for 6 days using protocols by Okae et al. or Karakis et al., staining for SDC1 and hCG. Primary CTBs are from the same placenta as those used for deriving T1 hTSCs in TUA medium.

(C) Confocal microscopy imaging of cells obtained during attempted derivation of hTSCs in TUC medium, differentiated to EVTs for 6 days using protocols by Okae et al. or Karakis et al., staining for NOTCH1 and HLA-G. Primary CTBs are from the same placenta as those used for deriving T1 hTSCs in TUA medium.

(D) Confocal microscopy imaging of cells obtained during attempted derivation of hTSCs in TUC medium, differentiated to STB for 6 days using protocols by Okae et al. or Karakis et al., staining for SDC1 and hCG. Primary CTBs are from the same placenta as those used for deriving T1 hTSCs in TUA medium.

Nuclei were stained with DAPI (blue). Scale bars are 100μm for all images.

### References

1. Karakis, V., Jabeen, M., Britt, J. W., Cordiner, A., Mischler, A., Li, F., San Miguel, A., and Rao, B. M. (2023) Laminin switches terminal differentiation fate of human trophoblast stem cells under chemically defined culture conditions. *Journal of Biological Chemistry*. 10.1016/j.jbc.2023.104650
